# Spatial Proximity Sequencing Maps Developmental Dynamics in the Germinal Center

**DOI:** 10.1101/2025.10.27.684659

**Authors:** Huili Wang, Junjie Xia, Peer Mohamed Suhail Mujibur Rahman, Bijentimala Keisham, Abinash Padhi, Yiyu Deng, Yan Li, Luke Vistain, Sarah Kim, Gabriel Mercado-Vasquez, Aly A. Khan, Marcus R. Clark, Savaş Tay

## Abstract

Spatial profiling of proteins and protein interactions is essential for immunology, signaling, development, and cancer. We present Spatial Proximity-Sequencing (Sprox-seq), a multi-omic technique that simultaneously measures proteins, protein complexes and mRNAs, where location of each molecule is also recorded. Sprox-seq profiled 32 proteins, 528 pairwise protein interactions and thousands of mRNAs across human tonsil tissues and germinal centers. Mapping protein interactions recapitulated RNA-defined tissue architecture in germinal centers, but also revealed much higher interaction complexity in the Light zone. Developmental trajectories inferred from protein interactions uncovered a B cell maturation pathway distinct from that inferred by RNA. Integrated protein-complex and mRNA analysis related spatially-enriched complexes with immune regulation and mitotic gene-expression pathways. Furthermore, Sprox-seq captured B cell-Follicular Dendritic Cell interactions mediated by the protein complex VLA-4–VCAM1 in the Light zone. Sprox-seq provides a multi-modal view of cell states and a powerful tool for studying protein and cellular interactions across tissues.

## Introduction

Spatially resolved measurement of key molecules in tissues has paved the way for understanding basic and disease phenotypes across many domains in biology, and may lead to the development and training of computational models of tissue organization and function^1–4^. Spatial transcriptomics has become popular among highly multiplexed techniques due to easy access to transcripts by existing genomic methods, and has mapped gene expression across tissue sections with high resolution and revealed new cellular identities^1,4–6^. However, transcriptomic data alone cannot capture many complex phenotypes and signaling events that govern cells and tissues. Proteins are the fundamental executors of cellular functions such as antigen recognition^7^, co-stimulation^8^, cell activation^9^, cell communication^10^ and effector function^11^. Proteins function through their dynamic expression, spatial localization, and especially through the formation of protein complexes both within and between cells. Thus, integrating transcriptomic, proteomic, and interactome (protein interaction) data with spatial resolution is critical for comprehensively characterizing cell states, studying functional tissue compartments, recording signaling dynamics, and understanding cell functions in intact tissues^12–16^.

Measurement of proteins with spatial resolution have traditionally relied on fluorescence microscopy^17^, including multiplexed methods such as Immuno-SABER^18^ and CODEX^19^ that enable the measurement of many protein species simultaneously. While preserving spatial information and providing subcellular resolution, these methods require iterative chemical staining and signal removal cycles that restrict throughput, risks signal degradation, and increases processing time^18,19^. Other techniques, such as SPOTS and DBiT combined with Cite-seq, use antibody–oligonucleotide conjugates (AOCs) with spatial transcriptomics to co-profile mRNAs and proteins with spatial resolution^3,20^. However, none of the above mentioned methods can detect protein interactions and complexes in situ.

Proximity ligation–based technologies have the potential to address these limitations. Methods such as Prox-seq^16^, Molecular Pixelation^12^, and iSeq-PLA^21^ combined AOCs and proximity ligation assay to detect protein expression and their interactions by converting them into DNA products. Sequencing-based platforms like Prox-seq and Molecular Pixelation quantify protein abundance and interactions by amplifying and sequencing the ligated products. However, both methods have been optimized for dissociated single cells and lack spatial resolution over tissues. Moreover, Molecular Pixelation does not provide transcriptomic readouts. iSeq-PLA preserves spatial context in intact tissue using rolling circle amplification and in situ fluorescence, but its reliance on iterative signal removal limits multiplexing (i.e. the number of molecular species that can be simultaneously measured), and the absence of mRNA detection restricts integrative analysis.

To overcome these limitations, we developed Spatial Proximity-Sequencing (Sprox-seq), a scalable experimental and computational platform that robustly integrates proximity ligation assay with spatial transcriptomics (Figures 1A and 1B). We first developed and validated a large panel of antibody-oligonucleotide conjugates (AOCs) suitable for proximity ligation assays in intact human tissues and enable multiplexed readout by existing spatial transcriptomics pipelines. We optimized several experimental conditions including tissue fixation and permeabilization, AOCs staining, and reduction of non-specific interactions. We also developed statistical/computational methods to remove background noise typically seen in proximity ligation assay-based readouts, which enabled quantification of protein complexes. We integrated gene expression with our proteomic readouts. As a result, Sprox-seq enables simultaneous in situ profiling of a large number of surface proteins, protein complexes and mRNAs in intact 2D tissue sections, preserving native tissue structure while supporting high multiplexed molecular readouts. Sprox-seq shows quadratic scaling of the number of protein interactions measured. The number of protein dimers measured in Sprox-seq scale quadratically with the number of proteins being targeted, which massively increases its multiplexing ability for high-content analysis of tissues.

**Figure 1.**
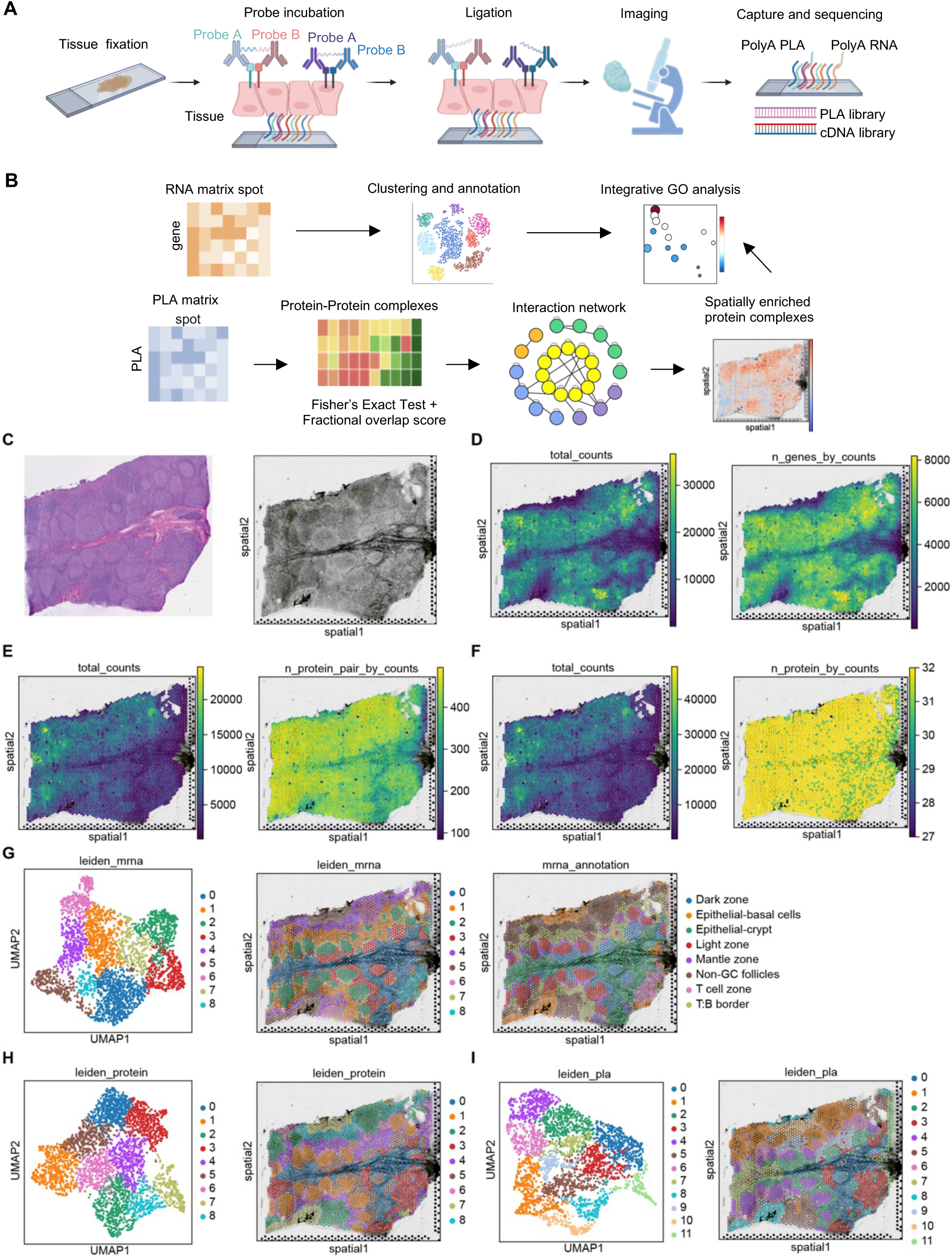
Spatial Prox-seq workflow and clustering analysis of human tonsil tissues. (A) Overview of the Sprox-seq experimental workflow. Human tonsil tissue sections are fixed onto Visium slides and incubated with AOC pairs. Upon spatial proximity, the probes are ligated and treated with USER enzyme to expose a poly (A) tail, enabling co-capture of PLA products and mRNAs via spatial barcodes. (B) Computational pipeline for RNA and PLA data analysis. RNA expression is analyzed using standard spatial transcriptomics workflows. PLA UMI counts are processed to infer protein expression and identify protein complexes using Fisher’s Exact Test. Protein fractional overlap analysis quantifies interaction strength. Cluster-specific interaction network was constructed using protein complexes identified within each cluster. Integration of RNA data enables functional annotation of spatially enriched protein interactions. Part of panel A and B were created using Biorender. (C) Histological reference and imaging. H&E staining of an adjacent tissue section (left) and brightfield image of the Visium slide used for capture (right). (D) Total mRNA UMI counts and number of detected genes per spot for sample A1. (E) Total PLA UMI counts and number of detected PLA pairs per spot for sample A1. (F) Total protein UMI counts and number of detected proteins per spot for sample A1. (G) UMAP plot of the clustering analysis based on RNA data (left), spatial distribution of the clusters (middle) and cluster annotation based on H&E morphology and RNA expression (right). (H) UMAP plot of the clustering analysis based on normalized protein data (left) and spatial distribution of the clusters (right). Protein data was derived from PLA UMI counts. (I) UMAP plot of the clustering analysis based on normalized PLA data (left) and spatial distribution of the clusters (right). See also Figures S1-S4 and Tables S1-S4.

We applied Sprox-seq to tissues of the human tonsil, an important secondary lymphoid organ that is responsible for fighting against pathogen infection^22,23^. Tonsil tissue is complex and contains multiple structurally distinct spatial regions, including primary follicles, mantle zone, germinal centers, T:B border zones, and epithelial regions. Various cells of the immune system such as B cells, T cells, and follicular dendritic cells (FDCs) interact in a spatially and temporally dynamic manner in tonsil tissue^22–25^. Germinal Centers, as the key functional structure of tonsil tissues, divided into Light Zone and Dark Zone, are the sites where antigen-activated B cells undergo somatic hypermutation and positive selection to generate high affinity antibodies. This process is tightly regulated by dynamic interactions between different cell types^24,26,27^. Tonsil tissue is an ideal test case for demonstrating the utility of Sprox-seq due to its highly compartmentalized and multi-cellular nature, and due to its reliance on protein interactions and cell–cell interactions across multiple spatial and temporal scales.

By using Sprox-seq, we spatially profiled 32 proteins and 528 pairwise protein interactions alongside thousands of mRNAs in tonsil tissue. Each spatial pixel revealed co-localized transcript, protein and protein-complex information across tissues, and multi-modal molecular maps of tonsil tissues have been generated. Clustering based on protein interaction data recapitulates the tonsil tissue structure similar to that defined by RNA. However, we have found important differences through protein complex-based analyses. Protein interaction networks within each spatial cluster revealed that Light zone demonstrates a much higher interaction network complexity compared to Non-GC follicles and Dark zone, consistent with its role as the sites where B cell interact with other cell types for selection during the maturation process.

Moreover, developmental trajectory analysis based on protein interactions revealed a maturation pathway spanning from Non-GC follicles to Dark zone and then to Light zone, which is distinct from that inferred by RNA data, offering a unique insight to understand B cell state transition. Twenty-four protein heterodimers were identified across all biological replicates and mapped along the pseudotime trajectory, including canonical protein complexes CD19–CD21, CD21–CD35, CD81–CD9, ITGA4–CD29 and several noncanonical interactions such as CD32–CD38, CD147–CD38, and CD20–CD32. Protein complexes CD19–CD21 and CD21–CD35 showed higher interaction strength in Non-GC follicles and Germinal Centers respectively, consistent with their roles in B cell activation and maturation.

Mapping protein expression along the same trajectory revealed that CD19 is more enriched in Non-GC follicles, whereas CD21 is more abundantly expressed in Light zone. The expression patterns of these proteins could not predict spatial enrichment of the corresponding complex CD19 – CD21 in Non-GC follicles. This indicated that protein interactions cannot be easily inferred from protein expression data alone, and highlighted the strength of direct protein-complex measurements such as that provided by Sprox-seq. Furthermore, by integrating transcriptomics data, we explored the functional implication of spatially enriched protein complexes. Pathway enrichment analysis of upregulated genes in spots with elevated protein complex interaction revealed the association of CD21–CD35 to cell division pathways in Germinal Centers and CD20–CD32 to immune activation in Non-GC follicles.

An advantage of Sprox-seq is its ability to directly capture cell – cell interactions across tissues using protein interactions between different cell types (in trans), which is distinct from cell-cell communication inferred from RNA co-expression. Single cell-based methods are typically unable to directly obtain cell – cell interaction information, as dissociation of single cells disrupts cell contacts. With our unique ability to measure protein complexes in tissues, we directly detected VLA-4 – VCAM1 complex mediated B cell-Follicular Dendritic Cell (FDC) interactions in Light zone of Germinal Centers. These analyses demonstrated the utility of our technology to detect cell– cell communication events across complex tissues.

## Results

### Development of experimental and computational pipelines of Spatial Prox-seq

The experimental and computational workflows of Sprox-seq are illustrated in Figures 1A and 1B, with detailed protocols provided in the Methods section. Briefly, 10-μm-thick human tonsil tissue sections were mounted onto spatial-transcriptomics (10x Visium) slides and fixed with 1% paraformaldehyde (PFA). A cocktail of AOCs pairs was applied to the tissue. Each protein was targeted with a pair of probes (Probe A and Probe B) generated by conjugating two distinct DNA oligonucleotides with same antibody, each probe carrying a unique DNA barcode that enabled identification of the corresponding protein. Upon spatial proximity, these probes were ligated to generate PLA products. To enhance capture efficiency, Probe B was modified with six terminal Uridine-5′-triphosphate (UTP). Following ligation, USER enzyme digestion removed the UTPs to release the poly(A) tail on the PLA products. After imaging to record tissue morphology, the tissue was permeabilized, allowing both poly(A)-tailed mRNAs and PLA products to be simultaneously captured by spatial barcodes on the Visium slide.

Following sequencing, both digital RNA and PLA matrices were generated. RNA data were analyzed following standard spatial transcriptomics pipelines and clustering results were manually annotated based on HE staining and typical marker expression. PLA products were recorded in the format {ProbeA}:{ProbeB}, where ProbeA and ProbeB represent the identities of the two antibodies ligated. Bidirectional PLA pairs (e.g., CD19A:CD21B and CD21A:CD19B) were merged into unique protein pairs (e.g., CD19–CD21) for the relative quantification. For PLA-derived protein quantification, UMI counts from Probe A and B targeting the same protein were summed to obtain the total expression per protein.

Proximity ligation can capture all targeted proteins colocalized within 50∼70 nm^16,28^. To identify functional protein pairs, we developed a modified statistical analysis pipeline to filter out proximity noise from PLA data (i.e. ligation of non-interacting protein pairs). This approach has been previously used to filter out non-specific PLA interactions in single cell readouts by proximity sequencing^16,29^. First, Fisher’s Exact Test^30^ was applied to PLA pairs to identify significant interactions. Pairs passing Fisher’s Exact Test were defined as protein complexes and further evaluated using protein fractional overlap analysis to quantify interaction strength (see Methods). This metric is adjusted from quantitative colocalization analyses in fluorescence microscopy, where it is used to evaluate the co-occurrence and correlation of labeled molecules^31,32^. The protein fractional overlap score provides a continuous, abundance-aware measure of interaction intensity across tissue spots. This two-step computational framework combines statistical significance testing and quantitative evaluation, enabling identification and quantification of protein complexes. After analysis with our pipeline, cluster-specific interaction network can be constructed based on the protein complexes identified within each cluster. Spatially enriched protein complexes were then analyzed across tissue regions and integrated with RNA expression data, enabling their functional interpretation.

### Simultaneous measurement of proteins, protein complexes and mRNAs across tissues

For use in human tonsil tissue, we developed a cocktail of 32 AOCs pairs, including 29 targeting specific proteins and 3 isotype controls (Tables S1 and S2). All primary antibodies were validated using immunofluorescence (IF) staining and the specificity of the AOCs was subsequently detected. Staining patterns produced by AOCs were found to closely match those obtained with the validated primary antibodies, indicating that oligonucleotide conjugation did not affect antibody specificity (Figure S1A).

The assay was performed on fresh-frozen human tonsil tissue sections, with three biological replicates: two spatially distinct regions from the same donor (samples A1 and B1) and one section from a different donor (sample D1). Both mRNAs and PLA products were simultaneously captured using the Sprox-seq workflow (Figures 1C–1E, S2A–S2C and S2E–S2G). Protein abundance data were generated from PLA UMI counts (Figures 1F, S2D and S2H).

At the median level, each spatial spot yielded 14,930 RNA UMIs spanning 5,403 genes, 4,799 PLA UMIs corresponding to 425 unique PLA pairs and 9,713 protein UMIs corresponding to 32 proteins across three replicates, demonstrating efficient co-detection of transcriptomic, proteomic and protein proximity features (Table S3). Furthermore, the three biological replicates showed high reproducibility across RNA, protein, and PLA measurements. Samples A1 and B1, derived from the same donor, exhibited slightly higher correlation values. These results highlight the robustness of our technology and support combinatorial analysis across all three samples (Figures S3A–S3C).

### Clustering analysis by Sprox-seq accurately identifies physiological zones and cell types within tonsil tissue

Unsupervised clustering based on RNA expression was performed for sample A1 and revealed nine major spatial tissue clusters, annotated based on HE staining and typical marker expression as Epithelial-basal cells, Epithelial-crypt, Light zone, Dark zone, Mantle zone, Non-GC follicles, T cell zone and T:B border^22^ (Figure 1G). PLA data and protein data were normalized using centered log ratio (CLR) methods following the approach used in Seurat. Clustering based on normalized protein abundance showed strong similarity with RNA-based clusters, confirming that our antibody panel captures the major cell populations in tonsil tissue (Figure 1H). Furthermore, clustering based on normalized PLA data, which captured spatial proximity information, also recapitulated similar tissue organization (Figure 1I). Similar results were observed in samples B1 and D1 (Figures S4A–S4F).

To validate whether Sprox-seq accurately captured protein expression patterns, IF staining of typical markers was performed on adjacent tissue sections using a combination of Probe A and Probe B. The spatial expression profiles of key markers, including B cell– associated markers (CD21, CD35), immunoglobulin isotypes (IgD, IgG), and T cell–specific markers (CD3, CD4), were observed to be highly consistent between IF and Sprox-seq (Figure 2A). Notably, the mouse isotype controls (mIgG1, mIgG2a, mIgG2b) showed lower signal, demonstrating negligible background binding (Figure S5A). Together, these results demonstrate that PLA-derived protein data with our cocktail reliably reflect both protein abundance and spatial localization within the tissue context.

**Figure 2.**
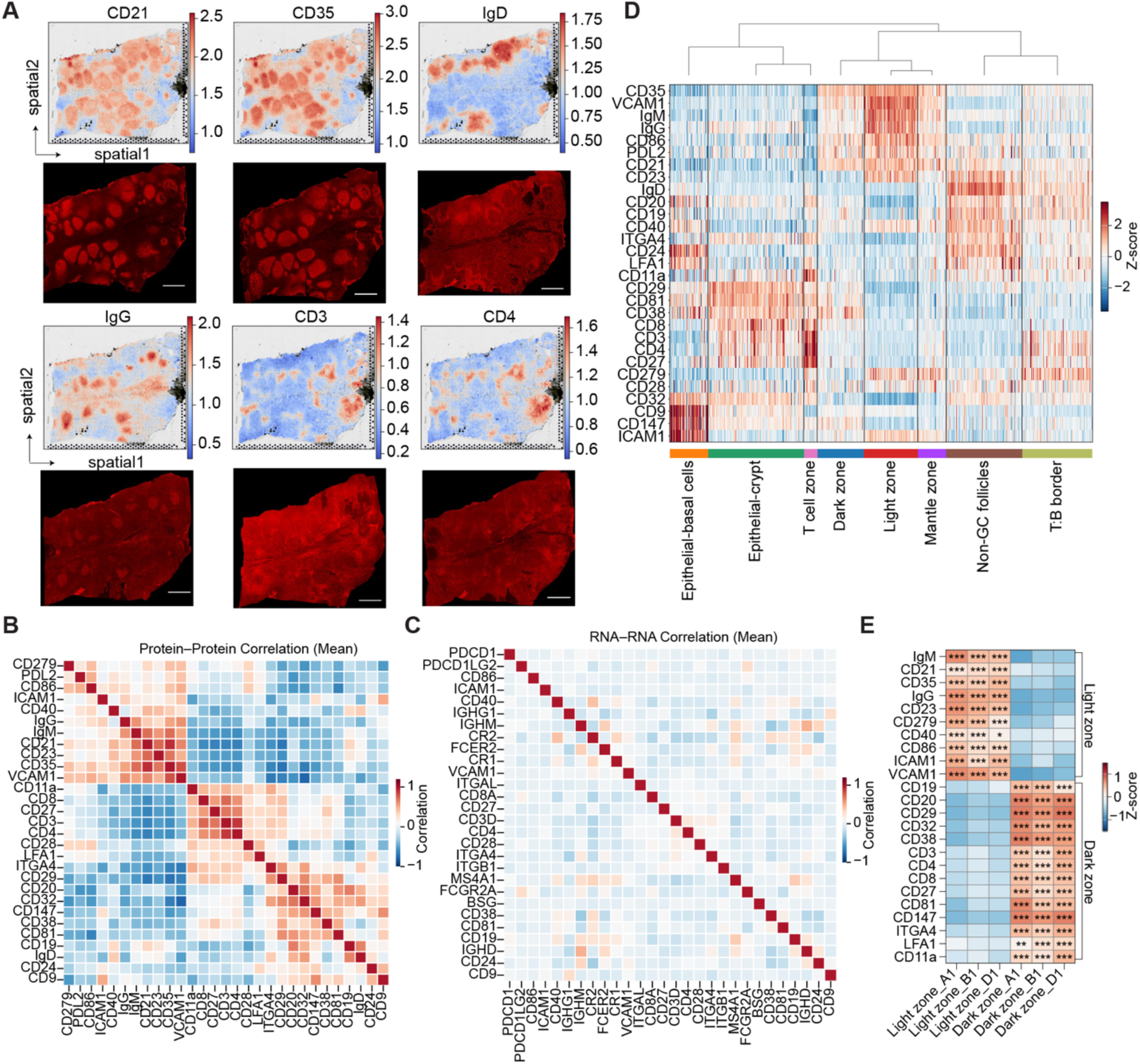
Validation and analysis of protein expression across spatial clusters in human tonsils. (A) Comparison of normalized protein expression measured by Sprox-seq (top), and IF staining from an adjacent tissue section using a mixture of Probe A and Probe B (bottom). B cell– associated markers (CD21, CD35), immunoglobulin isotypes ( IgD, IgG) and T cell markers (CD3, CD4) are shwon. Scale bar: 1mm. (B–C) Average Pearson correlation values among 29 proteins (B) and their corresponding genes (C) across three biological replicates. The correlation was calculated using normalized expression values. (D) Heatmap showing protein expression across spatial clusters. Rows represent proteins, and columns representing spots are grouped by clusters annotated based on RNA data. Normalized expression values were scaled to visualize relative protein abundance across regions. (E) Heatmap showing protein expression differences between Dark zone and Light zone across three biological replicates. Normalized expression values were scaled within each sample. Statistical significance for panel E was assessed using the two-sided Mann–Whitney U test with adjusted *P* values calculated by the Benjamini–Hochberg method, and significant differences are annotated with asterisks (* *P* <0.05, ** *P* <0.01, *** *P* <0.001). See also Figures S5-S8.

These analyses resulted in simultaneous measurement of protein abundance, protein complexes and mRNAs across ∼3000 spots for each tonsil tissue specimen we analyzed (Figures 1C–1F and S2A–S2H). The results created 2D spatial maps displaying the expressions of these functional molecules across tissues, resulting in visualization and molecular annotation of the tissues in a multi-dimensional manner. These maps and subsequent clustering analysis enabled identification of canonical cell types and protein species that exist in human tonsil tissue.

### Protein correlation analysis and differential expression analysis across spatial clusters

Next, protein co-expression patterns were analyzed, and the average pairwise correlations across three samples were calculated (Figure 2B). Four distinct modules were observed. Proteins CD21, IgM, CD23, IgG, CD35, and VCAM1 clustered together. A strong correlation pattern was also observed among co-signaling molecules such as CD279, PDL2 and CD86. T cell markers CD3, CD4, CD8 and CD27 formed a correlated module, and another module consists of CD147, CD38, CD29, CD81, CD20 and CD32. In contrast, similar correlation analysis performed on RNA expression of same genes showed much weaker patterns (Figure 2C), further demonstrating the importance of measuring protein signals.

Protein expression was compared across spatial clusters in sample A1, with annotations based on RNA data (Figure 2D). The T cell module, including CD3, CD4 and CD27, was mainly localized to T cell zone. Moderate expression of T cell markers CD3, CD4, CD279 and naïve B cell markers CD19, CD20 and IgD were observed in T:B border, indicating the coexistence of T cells and B cells in this region. IgD, CD19, CD20, CD21, CD23, CD24 and CD40 were expressed in Non-GC follicles, consistent with the existence of mature naïve B cell state in this region. CD9, CD147 and ICAM1 were enriched in Epithelial-basal cells, while CD29 and CD81 were observed in Epithelial-crypt regions. Light zone and Dark zone shared similar expression profiles but showed difference in expression levels. Similar spatial expression patterns were observed in samples B1 and D1, particularly in B1. Sample D1 showed some variations in epithelial regions and lacked Non-GC follicles, suggesting distinct physiological states among the tonsil samples (Figures S6A and S6B).

To more precisely compare protein expression profiles between Light zone and Dark zone, data from all three samples were jointly analyzed. Clustering and annotation of Dark zone and Light zone were confirmed by the expression of canonical markers (CD83, LMO2, BCL2A1 for Light zone; CXCR4, AICDA, MME and FOXP1 for Dark zone) and the enrichment of FDC markers (FDCSP, CLU, CR2, CXCL13) and T follicular helper markers (Tfh) (PASK, TIGIT, IFITM1) in Light zone^27,33^ (Figures S7A and S7B). Protein expression analysis revealed that proteins IgM, CD21, CD35, IgG, CD23, CD279, CD40, CD86, ICAM1 and VCAM1 were significantly enriched in Light zone (Figure 2E), consistent with its established role as a region of B cell-FDC interactions and B cell maturation^26,27,34^. In contrast, other proteins were more highly expressed in Dark zone (Figure 2E).

Finally, the correlation between RNA and protein expression levels across the panel was analyzed across three samples. Moderate to strong correlations were observed for proteins such as CD19, CD20, CD21, CD35, VCAM1, CD3, CD40, IgD, IgM and IgG, while weaker correlations were detected for others (Figure S8A), consistent with previous observations of gene-dependent variability between RNA and protein levels^35,36^.

Taken together, these findings demonstrate that our PLA-derived protein data and data analysis pipeline provide reliable measurements of protein expression, and offer an additional layer of biological insight.

### Spatial profiling of protein complexes across tonsil sub-regions reveals a positive association between network complexity and B cell maturation

To systematically identify significant protein complexes from raw PLA counts (Figures 3A, S9A and S10A), Fisher’s Exact Test was applied and co-binding events with adjusted p-values below 0.05 in the test were considered as significant. The spatial prevalence of each protein complex was further summarized as the ratio of positive spots, defined as the proportion of tissue spots where the complex was detected as statistically significant (see Methods).

**Figure 3.**
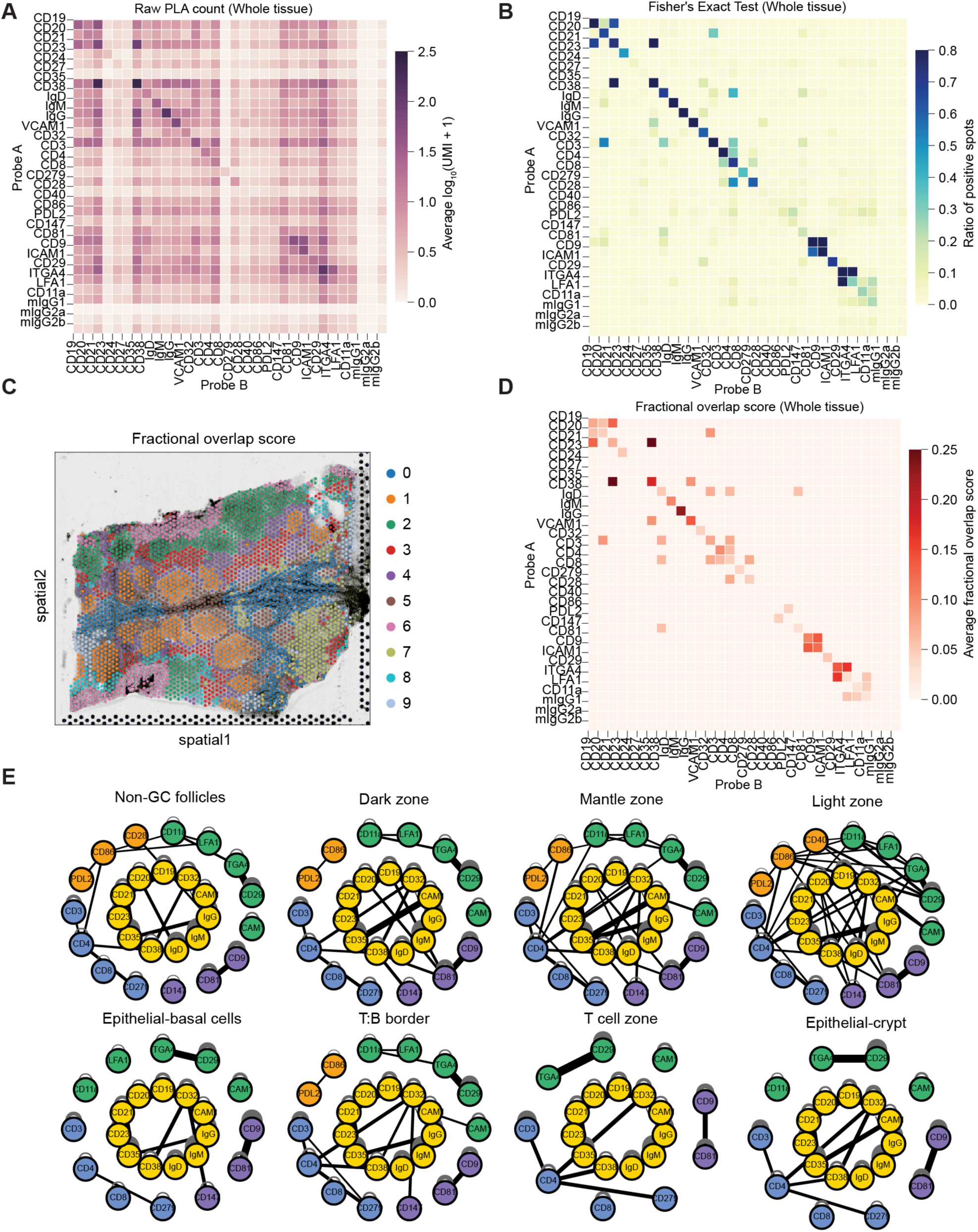
Spatial profiling of protein complexes across tonsil sub-regions reveals a positive association between network complexity and B cell maturation. (A) Heatmap showing raw PLA counts (log-transformed) for all detected PLA pairs across sample A1, as measured by Sprox-seq. (B) Heatmap of PLA pairs after Fisher’s Exact Test (adjusted *P* < 0.05) across sample A1, indicating enriched protein complexes. Values represent the ratio of positive spots. (C) Spatial plot of clustering analysis based on protein fraction overlap scores of protein pairs. (D) Heatmap of protein fractional overlap scores for protein pairs across sample A1. Only protein pairs with a ratio of positive spots greater than 0.05 are shown. (E) Protein interaction networks constructed for each spatial cluster. Nodes represent proteins and are colored by associated cell type or functional category: gold, B cell; blue, T cell; orange, co-signaling molecules; green, integrins; purple, tetraspanins. Edges denote protein complexes, with thickness proportional to the average protein fractional overlap score within the cluster. Only protein pairs with a positive spot ratio higher than 0.15 within the given cluster are displayed. See also Figures S9 and S10.

Analysis at the whole-tissue level revealed numerous canonical protein complexes, which are known to exist in human tonsil (Figures 3B, S9B and S10B). Many homodimers were consistently detected with high ratio of positive spots. Canonical heterodimers, including CD19–CD21^37^, CD21–CD35^38^, CD81–CD9^39^, and CD29–ITGA4^40^, were identified and all of them exhibited high positive spot ratios, suggesting their highly stable interactions. Several noncanonical interactions, including CD32–CD38, CD147–CD38, and CD20–CD32, were also identified, suggesting their interactions in membrane microdomains. Most PLA pairs involving mouse isotype controls failed to pass the statistical test, further supporting the robustness of the analytical framework (Figures 3B, S9B and S10B). The finding of these canonical protein complexes in tonsil tissue supports the accuracy of our technology in identifying protein complexes.

Fractional overlap scores were then computed for each protein pair to quantify their interaction strength. Clustering based on protein fractional overlap scores recapitulated spatial tissue patterns consistent with those obtained from transcriptomic data, indicating that the protein interaction strength also captures tissue organization (Figures 3C, S9C, and S10C). Among protein pairs passing Fisher’s Exact Test, those with higher ratio of positive spots also exhibited greater fractional overlap scores, indicating a positive association between spatial prevalence and interaction strength (Figures 3D, S9D and S10D).

To characterize the global interaction landscape, protein interaction networks were constructed for protein complexes with the ratio of positive spots higher than 0.15 within each spatial cluster. In these networks, node represents proteins, edge thickness indicates the average protein fractional overlap score within the cluster, and node color reflects the associated cell type or functional category. Cluster-featured interaction networks revealed region-specific interaction patterns (Figures 3E, S9E and S10E). B cell–enriched regions, including Non-GC follicles, Dark zone, Light zone and Mantle zone showed greater interaction complexity compared to epithelial regions and T cell zone, reflecting the design of the antibody panel. T:B border displayed a more connected network than T cell zone, which may suggest the presence of T–B cell crosstalk in this region.

Among the B cell–enriched regions, Non-GC follicles displayed the lowest level of interaction, followed by Dark zone and Mantle zone, with Light zone showing a particularly enriched pattern of intercellular connections, characterized by interactions crossing proteins from distinct immune cell types and different functional categories. This pattern demonstrates a positive association between interaction network complexity and B cell maturation state across these clusters. The elevated network complexity observed in Light zone may reflect its role in mediating B cell selection through dynamic communication with other cell types^27,33,41^. In contrast, Dark zone exhibited a less integrated network, indicating a less interactive microenvironment (Figures 3E, S9E and S10E). Some differences were observed for interaction networks among biological replicates (Figures 3E, S9E and S10E), revealing dynamic changes of interaction profiles resulted from heterogeneity of tissue states. Notably, the positive association between network complexity and B cell maturation within B–cell enriched clusters was consistently observed across all three samples.

Taken together, our analytical pipeline allowed the quantification of protein complexes and the construction of spatially resolved protein interaction networks, revealing coordinated communication within and between cell types across tissue regions. These findings emphasize the value of protein interaction profiling using Sprox-seq, which captures intercellular communication dynamics that are not detectable at the transcriptomic or protein expression level.

### Trajectory analysis based on protein interaction strength reveals a maturation pathway within B cell–enriched clusters, distinct from that inferred from RNA

To investigate dynamic changes of protein interactions during B cell maturation, B cell–enriched clusters from three samples were jointly analyzed. Regions representing more naïve B cell states (Non-GC follicles from samples A1 and B1), and more mature Germinal Center states (Light zone and Dark zone from samples A1, B1 and D1) were combined for further analysis after batch effect correction using Harmony^42^ (Figure S11A). After batch effect removal, clustering was performed based on both protein fractional overlap scores and transcriptomics profiles (Figures 4A and 4B).

**Figure 4.**
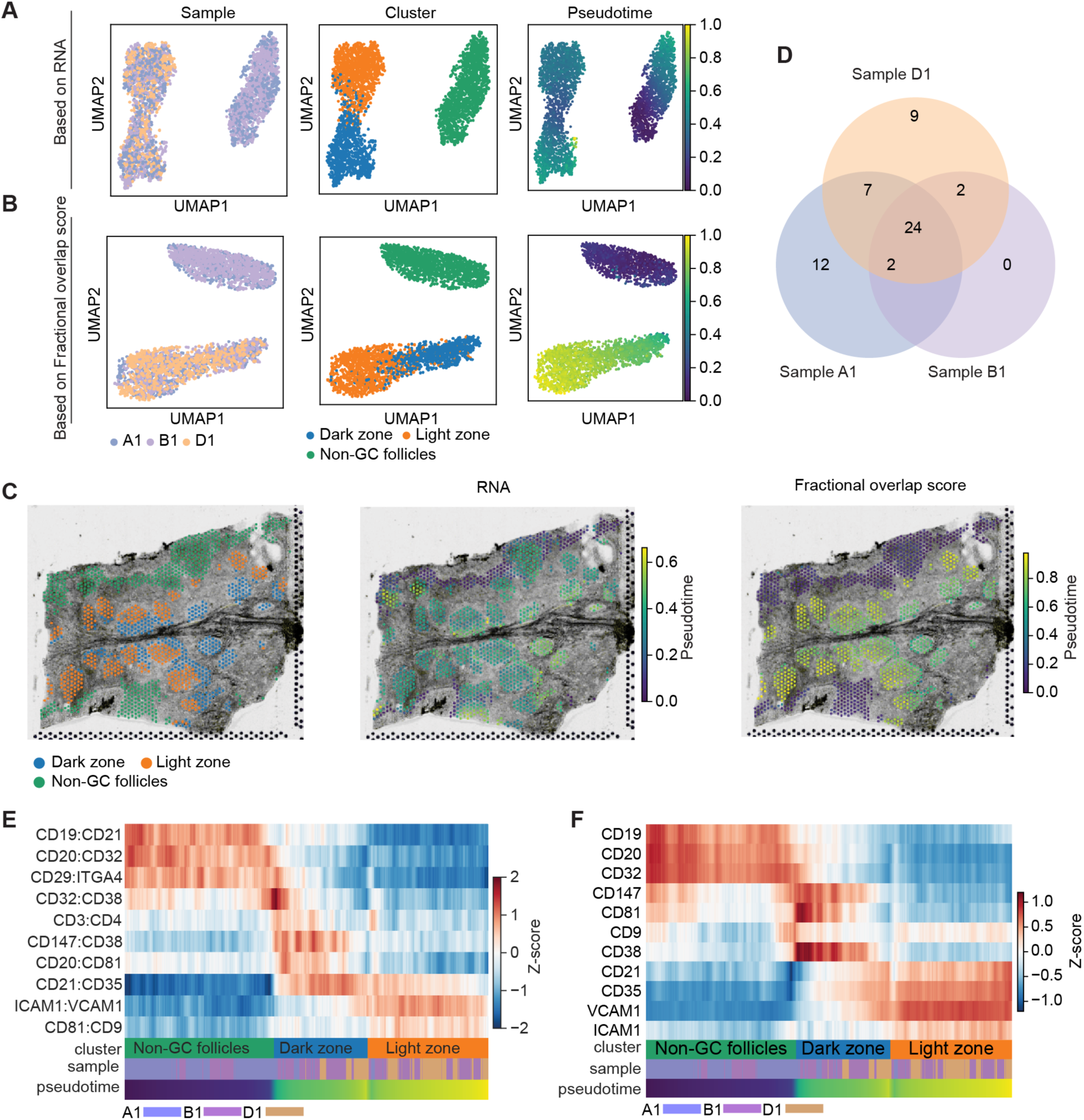
Trajectory analysis based on interaction strength reveals a maturation path distinct from that inferred from RNA within B cell–enriched clusters. (A–B) Pseudotime trajectory analysis using diffusion pseudotime (DPT) inferred from RNA data (A) and protein fractional overlap scores (B) of all three samples after batch correction with Harmony; the trajectory was rooted in Non-GC follicles. Left: UMAP plot showing spots colored by samples; Middle: UMAP plot colored by clusters based on RNA annotations; Right: UMAP plot showing pseudotime progression. (C) Spatial plot of selected clusters based on RNA annotations, including Dark zone, Light zone and Non-GC follicles (left), spatial plot of pseudotime values inferred from RNA expression (middle), and spatial map of pseudotime progression inferred from protein fractional overlap scores of protein pairs (right). (D) Venn diagrams showing the number of heterodimers consistently detected across three biological replicates. Protein pairs were defined as consistent if they passed Fisher’s Exact Test in all three replicates and exhibited a positive spot ratio greater than 0.15 in at least one of the selected clusters within each sample. (E) Heatmap showing scaled protein fractional overlap scores of selected heterodimers along the pseudotime trajectory. Spots are ordered based on pseudotime values and grouped by clusters. Z-score normalization was applied to each protein pair across spots from selected clusters. (F) Heatmap showing scaled expression levels of selected proteins along the same pseudotime trajectory. Normalized protein expression values were scaled within each protein. See also Figure S11.

Pseudotime trajectories were inferred from both transcriptomic profiles and protein fractional overlap scores using diffusion pseudotime (DPT)^43^, revealing two spatiotemporal patterns of B cell maturation (Figures 4A–4C). Based on RNA expression, a trajectory was inferred within Non-GC follicles cluster, uncovering two major subpopulations that reflect slightly different activation states of naïve B cells (Figures S11B and S11C). In this model, Dark zone and Light zone shared similar maturation degree, with Dark zone exhibiting a slightly higher maturation state, potentially suggesting increased transcriptional activity. In contrast, the trajectory obtained from protein fractional overlap scores presented a different progression path, starting from Non-GC follicles and extending through Dark zone to Light zone. The two subpopulations defined by RNA pseudotime analysis did not show much difference to the trajectory based on interaction strength. However, a clear transition from Non-GC follicles to Germinal Center zones (Dark zone and Light zone) was observed, with Light zone demonstrating the highest degree of maturation, consistent with our finding of a positive association between network complexity and B cell maturation state within the clusters.

To characterize the dynamic changes in protein interactions during B cell maturation, the trajectory based on protein fractional overlap scores was selected as reference for downstream analysis. Protein pairs were retained only if they passed Fisher’s Exact Test and had the ratio of positive spots greater than 0.15 in at least one of the selected clusters within each sample. Following these criteria, 22 homodimeric and 24 heterodimeric pairs were consistently identified (Figures 4D and S11D).

Heterodimeric pairs were further aligned across clusters along the pseudotime trajectory. While most integrin-mediated protein interactions did not exhibit strong cluster specificity, a few heterodimers demonstrated distinct distribution patterns (Figures 4E and S11E). For example, CD19–CD21 and CD20–CD32 were specifically enriched in Non-GC follicles, whereas CD21–CD35 was more prevalent in Light zone and Dark zone. The distinct distributions of CD19–CD21 and CD21–CD35 are consistent with their established roles in B cell activation and B cell maturation^38,44–47^ respectively, further supporting the accuracy of our protein-complex readouts with Sprox-seq. Furthermore, the expression of proteins forming these heterodimers was also aligned along the trajectory (Figure 4F). CD19 was enriched in Non-GC follicles and CD21 was more abundantly expressed in Light zone. The enrichment patterns of these proteins did not always align with the interaction strength of their corresponding protein complexes. The spatial discordances between individual protein expression and their associated interactions reinforce that protein interaction patterns identified by Sprox-seq cannot be directly inferred from data obtained using technologies that only measure protein abundance^3,20^.

In addition to the identification of a protein-interaction driven B cell maturation pathway, these analyses showed the compatibility of Sprox-seq with existing analytical methods that infer cellular dynamics, and the strength of Sprox-seq in integrating transcriptomic data with protein interactions to generate new insights into the dynamics of cellular transitions.

### Sprox-seq integrates transcriptional and protein-interaction data and enables functional annotation of spatially enriched protein complexes

The spatial distribution patterns might indicate potential roles for protein complexes in tissue function. To explore this, we integrated protein complex readouts with transcriptomic data from the same spatial locations. The CD21–CD35 complex was significantly enriched in Dark zone and Light zone compared to Non-GC follicles in tonsil (Figures 5A – 5C). Differential gene expression analysis was performed between high and low CD21–CD35 interaction spots within Germinal Centers in sample A1, identifying 278 upregulated and 97 downregulated genes (adjusted *P* <0.05 and |log_2_ (fold change)| >0.5), indicating significant changes in RNA expression (Figure 5D). Multiple cell cycle regulators, including CENPF, PLK1, KIF20A, TTK, SPC24, NEK2 and H3C7 were observed among the top 20 upregulated genes (Figure 5E) and pathway enrichment analysis of all upregulated genes in high CD21– CD35 interaction spots revealed significant enrichment of E2F targets, G2M checkpoint, mitotic spindle, mTORC1 signaling and DNA repair pathways (Figure 5F), demonstrating a positive association between CD21–CD35 interaction and cell mitosis. Similarly, pathway enrichment analysis performed on sample B1 also revealed a positive association between high CD21–CD35 interaction with active cell proliferation state (Figures S12A–S12D).

**Figure 5.**
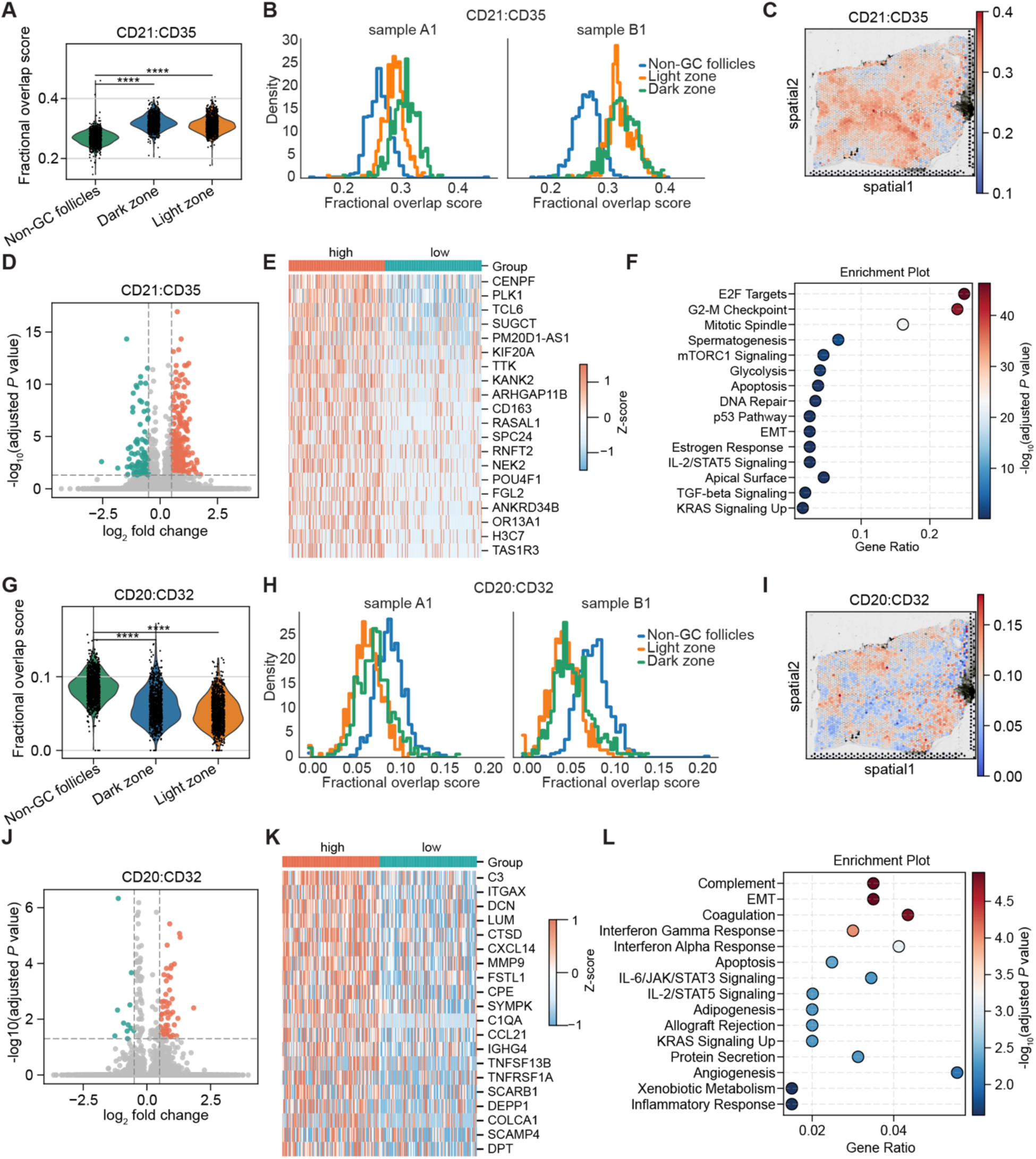
Sprox-seq enables the functional annotation of spatially enriched protein complexes. (A) Violin plot of raw protein fractional overlap scores for CD21– CD35 across selected clusters of three samples. Each dot represents a single spatial spot. (B) Histograms of CD21–CD35 raw protein fractional overlap scores across selected clusters from samples A1 and B1. (C) Spatial plot of CD21–CD35 raw protein fractional overlap scores in sample A1. (D) Volcano plot of differentially expressed genes between high and low CD21–CD35 interaction spots within Germinal Centers (Light zone and Dark zone) in sample A1. High and low groups were defined as the top 20% and bottom 20% of protein fractional overlap scores. (E) Heatmap showing the top 20 upregulated genes. (F) Hallmark pathway enrichment analysis (MSigDB Hallmark 2020) of 278 upregulated genes in high CD21–CD35 interaction group. (G) Violin plot of raw protein fractional overlap scores for CD20 – CD32 across selected clusters of three samples. Each dot represents a single spatial spot. (H) Histograms of CD20–CD32 raw protein fractional overlap scores across selected clusters from samples A1 and B1. (I) Spatial plot of CD20–CD32 raw protein fractional overlap scores in sample A1. (J) Volcano plot of differentially expressed genes between high and low CD20– CD32 spots within Non-GC follicles from samples A1 and B1. High and low groups were defined as the top 20% and bottom 20% of protein fractional overlap scores. (K) Heatmap showing the top 20 upregulated genes. (L) Hallmark pathway enrichment analysis (MSigDB Hallmark 2020) of 52 upregulated genes in high CD20–CD32 interaction group. Statistical significance for panels A and G was assessed using the two-sided Mann–Whitney U test; asterisks indicate significance levels: *P* < 0.0001 (****). Gene differential expression analsyis for panels D and J was analyzed using the Wilcoxon rank-sum test and adjusted *P* values were calculated by the Benjamini–Hochberg method. Upregulated genes (log_2_(fold change) > 0.5 and adjusted *P* < 0.05) are labeled as orange; Downregulated genes (log_2_(fold change) < -0.5 and adjusted *P* < 0.05) are labeled as teal. See also Figures S12 and S13.

CD20 – CD32 was significantly enriched in Non-GC follicles (Figure 5G– 5I). This interaction has not been previously measured directly, though recent literature supports the simultaneous upregulation of CD20 and CD32 during viral infection^48^ and the existence of an interaction between CD20 – CD32 mediated by Rituximab^49^. To explore the possible functional relevance of CD20–CD32 interaction, we performed differential gene expression analysis between high and low CD20–CD32 interaction spots within Non-GC follicles in sample A1, identifying 52 upregulated and 11 downregulated genes (Figures 5J). Among the top 20 upregulated genes, complement components (C3, C1QA), chemokines (CXCL14, CCL21), extracellular matrix molecules (DCN, LUM, MMP9, FSTL1, DPT and COLCA1) and the immune signaling receptor gene TNFRSF1A were observed (Figure 5K). Pathway enrichment analysis of all the upregulated genes in high CD20–CD32 interaction spots revealed significant enrichment of pathways associated with complement activation, extracellular matrix remodeling and interferon-γ response (Figure 5L). Consistent results were observed in sample B1 (Figures S12E – S12H). These suggest that CD20 – CD32 interaction on cells in Non-GC follicles may play a role in immune complex recognition, complement activation and the coordination of innate immune responses.

CD19–CD21 was also significantly enriched in Non-GC follicles (Figures S13A–S13D). However, differential gene expression analysis of high interactions revealed no significant changes in RNA expression, possibly because this interaction is distributed among the majority of naïve B cells, which primes B cells for activation.

Overall, our technology integrated transcriptional and protein-interaction data in a spatially resolved manner, and discovered associations between transcriptional programs and spatially enriched protein complexes from intact tonsil tissues. This integrated analysis enabled the identification of potentially new protein complexes defining distinct stages of B cell activation and maturation in immunological tissues.

### Sprox-seq captures VLA-4 – VCAM1 mediated interactions between B cells and Follicular Dendritic Cells

Sprox-seq does not require tissue dissociation, thereby preserving native tissue structure and maintaining intact cell–cell interactions. In our antibody panel, ITGA4 and CD29, two subunits of the integrin VLA-4, and VCAM1 form a well-known interaction between B cells and FDCs^25^. VLA-4 is highly expressed on activated T and B cells, while VCAM1 is predominantly expressed on FDCs. Across three biological replicates, VCAM1 was significantly enriched in Light zone (Figure 2E), whereas ITGA4 and CD29 showed a wide distribution outside of Germinal Centers (Figures 6A–6C). Due to their exclusive expression patterns in different cell types, the chance of cis interactions between these proteins is low. In our experiments, spatial analysis of both ITGA4–VCAM1 and CD29–VCAM1 interaction demonstrated highest enrichment in Light zone across all three samples, consistent with B cell–FDC interactions in this region (Figures 6D–6G, S14A and S14B).

**Figure 6.**
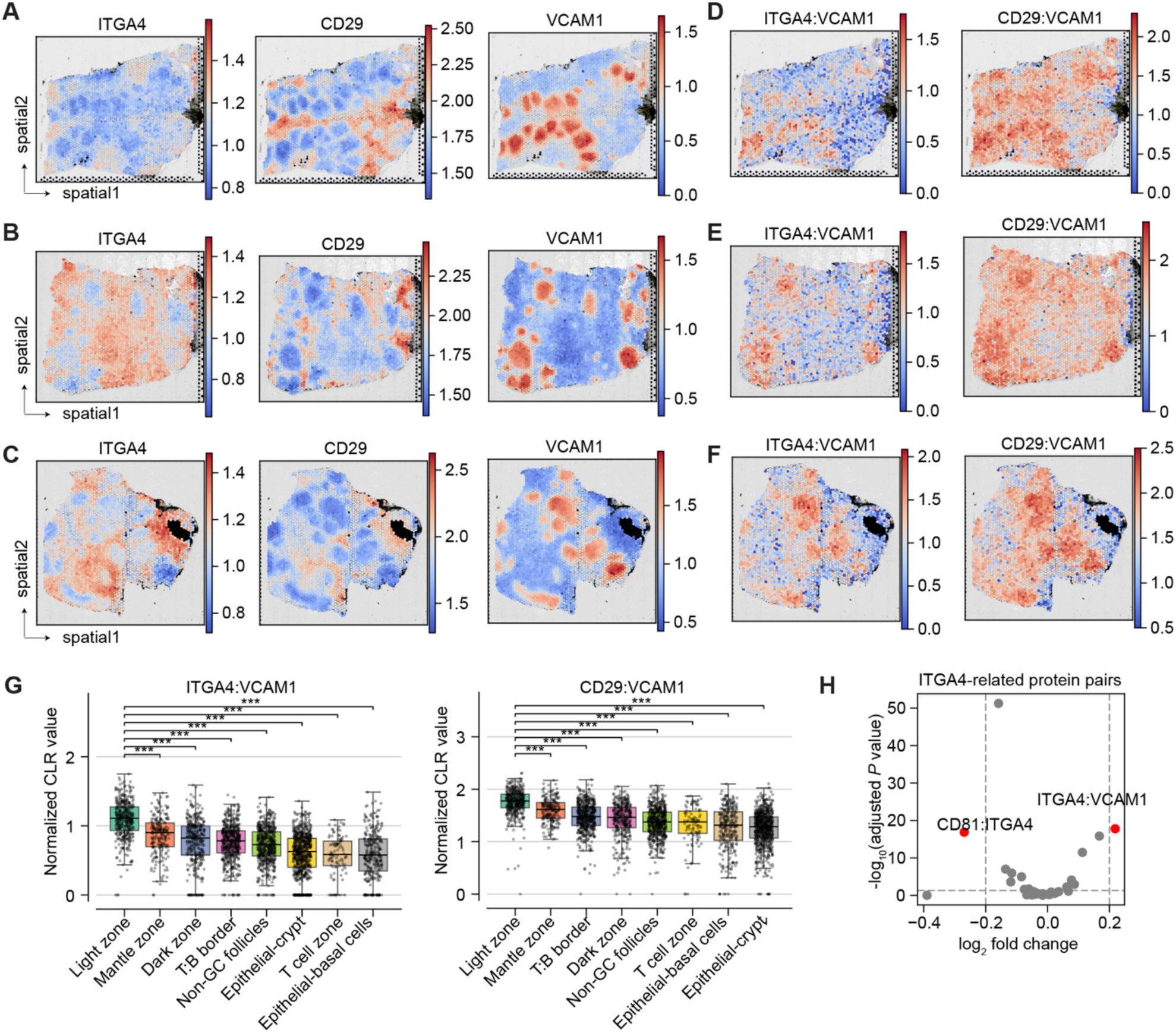
Sprox-seq captured VLA-4–VCAM1 mediated cell–cell interaction in germinal centers. (A-C) Spatial plot of normalized expression of the two VLA-4 subunits, ITGA4 (left) and CD29 (middle), and the corresponding ligand VCAM1 (right) across all three samples: samples A1 (A), B1 (B) and D1 (C). (D–F) Spatial plot of normalized value of protein pairs ITGA4–VCAM1 (left) and CD29–VCAM1 (right) across all three samples: samples A1 (D), B1 (E) and D1 (F). (G) Comparison of normalized value of ITGA4–VCAM1 (left) and CD29–VCAM1 (right) across all clusters in sample A1. (H) Volcano plot showing differential interactions of ITGA4-related protein pairs between Light zone and Dark zone. Each dot represents a protein pair involving ITGA4. Protein pairs with |log₂ fold change| > 0.2 and adjusted *P* < 0.05 are highlighted in red. Statistical significance for panels G and H was assessed using the two-sided Mann–Whitney U test and *P* values were adjusted by the Benjamini–Hochberg method; asterisks indicate significance levels: *P* < 0.001 (***). See also Figure S14.

To evaluate the interaction specificity of ITGA4 in Light zone, the interaction strength of all ITGA4-involved protein pairs was compared to the strength detected in Dark zone. ITGA4–VCAM1 interactions showed the highest difference, suggesting that ITGA4 had the strongest interaction specificity with VCAM1 in Light zone (Figure 6H). Taken together, these results demonstrated that Sprox-seq technique enables the direct capture of protein interactions mediating cell–cell communications in situ, across intact tissues.

## Discussion

Simultaneous spatial mapping of protein abundance, protein interactions and gene expression is essential for understanding the behavior of cells and their organization within tissues. We introduced Sprox-seq, a spatial multi-omic method that integrates proximity ligation assays with spatial transcriptomics to simultaneously profile mRNAs, surface protein abundance, and protein complexes within intact tissue sections. Application of Sprox-seq to human tonsils and especially germinal centers within them resulted in insights and findings in B cell physiology that could not be achieved by analysis of RNA or protein expression alone.

We comprehensively validated our Sprox-seq protein readouts using orthogonal methods like immunofluorescence, tissue staining and structural analysis. We also developed an analysis framework to enable robust identification of protein complexes and quantitative measurement of their strength. Our combined analyses captured the expected physiological tissue compartments, cell types and protein and protein-complex expression patterns. We used Sprox-seq to show that clustering based on either PLA-derived protein expression or protein interaction profiles can recapitulate transcriptome-defined tissue organizations.

Application of our method to human tonsil tissues revealed numerous homodimers and canonical (expected) heterodimers, such as CD19–CD21, CD21–CD35, CD29–ITGA4, and CD81–CD9, along with a few noncanonical and potentially novel complexes like CD32– CD38, CD147–CD38 and CD20–CD32. Cluster-specific protein interaction networks constructed from these protein complexes further revealed that network complexity is positively associated with B cell maturation state within the clusters. Particularly, a much denser interaction network was observed in the germinal center Light zone compared to Dark zone, consistent with the role of Light zone in mediating cell-cell communications during B cell maturation.

Focusing on B cell-enriched regions in the Germinal Center, we leveraged both RNA expression and protein interaction strength to reconstruct developmental trajectories covering from more naïve B cells (Non-GC follicles) to mature Germinal Center cells (including Light zone and Dark zone). Notably, the two trajectory analyses revealed distinct maturation paths. The trajectory based on protein interaction strength suggested a progression from naïve B cells to Dark zone and then to Light zone, highlighting that protein interaction offers unique insights into B cell state transitions, improving our understanding of B cell maturation.

To explore the functional associations of these protein complexes, 24 heterodimeric protein pairs significantly detected across biological replicates were aligned along the pseudotime trajectory. CD19–CD21, CD20–CD32 showed strong enrichment in Non-GC follicles, and CD21-CD35 showed higher enrichment in Germinal Centers. Furthermore, gene expression analysis of spots with high CD21–CD35 and CD20–CD32 interaction linked their patterns to the function. These results demonstrate that protein interaction profiles can reveal distinct immune cell states.

Distinct from inferring cell communications from RNA co-expression, Sprox-seq enables the direct capture of cell–cell interactions by detecting interacting protein pairs across different cell types. Indeed, VLA-4–VCAM1 protein complex mediated interaction between B cells and FDCs was observed in the Light zone of Germinal Centers across all three biological replicates.

Despite these advances, Sprox-seq has current limitations. Firstly, the quality and specificity of antibody-oligonucleotide conjugates is critical. Like other antibody-based methods, rigorous pre-validation of antibody panels and oligos is essential prior to Sprox-seq. The spatial resolution of the current results is limited by the size of Visium capture spots we used, which does not reach single-cell resolution. Incorporation of higher-resolution capture technologies ^5,50,51^ would further enhance the power of our method. Measurement of intracellular proteins and complexes is challenging due to limited access to intracellular proteins and higher background noise, and remain beyond the scope of this current study. While Sprox-seq provides a qualitative way of visualizing cell-cell interactions by capturing protein proximity signals between neighboring cells within a spot, it does not yet enable full quantification of such trans-interactions. Future statistical frameworks that integrate probabilistic modeling and spatial information would enable quantification of the detected intercellular interactions.

In conclusion, Sprox-seq provides a comprehensive and scalable platform for simultaneous in situ profiling of gene expression, protein abundance, and protein interactions across tissues. It enables the characterization of many known and potentially novel protein interaction networks across different spatial regions, as well as dynamic proteomic trajectories of cells. In addition to the identification of several immunological phenomena driven by protein-protein interactions, our results showed the compatibility of Sprox-seq with existing analytical methods that infer cellular dynamics, and its capability to integrate transcriptomics with protein expression and interactions to generate new insights into the dynamics of cellular transitions. While this study focused on human tonsil tissue and Germinal Centers within them, the technology can be applied to other tissues to explore developmental biology, neuronal biology, and disease progression.

## Resource availability

### Lead contact

Further information and requests for resources and reagents can be directed to Savaş Tay.

### Materials availability

All unique reagents used in this study are available upon request from the lead contact with the guidance of Materials Transfer Agreement of University of Chicago.

### Data And Code Availability

The raw data and count matrix have been deposited in the NCBI Gene Expression Omnibus under accession numbers GSE304749. The python files for reproducing data analysis are available at https://github.com/jjxia955/Spatial-Prox-seq.

## Acknowledgements

This work was supported by NIH grant AI175321 (S.T., M.R.C), P. Allen Distinguished Investigator Award (S.T), and Chan-Zuckerbeg Chicago Biohub Investigator Award (S.T. and M. R. C.). We acknowledge Human Tissue Resource Center, The University of Chicago Genomics Facility, The University of Chicago Cytometry and Antibody Technology facility, Integrated Light Microscopy Core, Cellular Screening Center and The University of Chicago Research Computing Center for their services. We thank Andrew Wang, Jonathan Matthews for discussions on data analysis. We thank Margaret Veselits for the help with sample treatment. We acknowledge the Center for Research Informatics Bioinformatics Core, University of Chicago, for support with code review and data analysis.

## Author contributions

H.W. and J.X. designed and performed the experiments and analyzed the sequencing data. P.M.S.M.R. and B.K. assisted with sequencing. A.P., Y.D., G.M.V, and S.K. contributed to antibody validation. H.W., J.X., and S.T. wrote the manuscript. M.R.C provided experimental guidance and participated in project design and data interpretation. L.V. contributed to data interpretation. Y.L. contributed to code review and data analysis. A.A.K. provided computational guidance. S.T. supervised the project. All authors reviewed and approved the final manuscript.

## Declaration of interests

The authors declare no competing interests.

## STAR★Methods

### Key Resources Table

| REAGENT or RESOURCE | SOURCE | IDENTIFIER |
| --- | --- | --- |
| **Antibodies** |  |  |
| Purified anti-human CD21 Antibody | BioLegend | Cat# 354902; RRID:AB_11219188 |
| Purified anti-human CD35 Antibody | BioLegend | Cat# 332402; RRID:AB_2890764 |
| Purified anti-human IgM Antibody | BioLegend | Cat# 314502; RRID:AB_493003 |
| Purified anti-human CD32 Antibody | BioLegend | Cat# 303201; RRID:AB_314333 |
| Purified anti-human CD19 Antibody | BioLegend | Cat# 302202; RRID:AB_314232 |
| Purified anti-human CD9 Antibody | BioLegend | Cat# 312102; RRID:AB_314907 |
| Purified anti-human CD81 (TAPA-1) Antibody | BioLegend | Cat# 349502; RRID:AB_10643417 |
| Purified anti-human CD38 Antibody | BioLegend | Cat# 356602; RRID:AB_2561794 |
| Purified anti-human IgG1 Antibody | BioLegend | Cat# 409904; RRID:AB_2650624 |
| Purified anti-human CD23 Antibody | BioLegend | Cat# 338502; RRID:AB_1279181 |
| Purified anti-human IgD Antibody | BioLegend | Cat# 348202; RRID:AB_10550095 |
| Purified anti-human CD3 Antibody | BioLegend | Cat# 300302; RRID:AB_314038 |
| Purified anti-human CD147 Antibody | BioLegend | Cat# 306202; RRID:AB_314586 |
| Purified anti-human CD20 Antibody | BioLegend | Cat# 302301; RRID:AB_314249 |
| Purified anti-human CD27 Antibody | BioLegend | Cat# 302802; RRID:AB_314294 |
| Purified anti-human CD54 Antibody | BioLegend | Cat# 353102; RRID:AB_11204426 |
| Purified anti-CD106 (VCAM-1) Antibody | BioLegend | Cat# 919801; RRID:AB_2565212 |
| Purified anti-human CD4 Antibody | BioLegend | Cat# 300502; RRID:AB_314070 |
| Purified anti-human CD8 Antibody | BioLegend | Cat# 344702; RRID:AB_1877104 |
| Purified anti-human CD28 Antibody | BioLegend | Cat# 302902; RRID:AB_314304 |
| Purified anti-human CD273 (B7-DC, PD-L2) Antibody | BioLegend | Cat# 329602; RRID:AB_1089010 |
| CD29 (Integrin beta 1) Monoclonal Antibody | Invitrogen | Cat# 14-0299-82; RRID:AB_1210468 |
| ITGA4 Monoclonal Antibody (HP2/1) | Invitrogen | Cat# MA1-80529; RRID:AB_928544 |
| Purified anti-human CD11a/CD18 (LFA-1) Antibody | BioLegend | Cat# 363402; RRID:AB_2564213 |
| CD11a/Integrin L/LFA-1 Chain Monoclonal Antibody (CRIS-3) | NeoBiotechnologies | Cat# 3683-MSM1-P1; RRID:N/A |
| Purified anti-human CD279 (PD-1) Antibody | BioLegend | Cat# 621602; RRID:AB_2820103 |
| Purified anti-human CD40 Antibody | BioLegend | Cat# 334302; RRID:AB_1236384 |
| Purified anti-human CD86 Antibody | BioLegend | Cat# 305402; RRID:AB_314522 |
| Purified anti-human CD24 Antibody | BioLegend | Cat# 311102; RRID:AB_314851 |
| Purified Mouse IgG1, $\kappa$ Isotype Ctrl Antibody | BioLegend | Cat# 401402; RRID:AB_2801451 |
| Purified Mouse IgG2a, $\kappa$ Isotype Ctrl Antibody | BioLegend | Cat# 401502; RRID:AB_2800437 |
| Purified Mouse IgG2b, $\kappa$ Isotype Ctrl Antibody | BioLegend | Cat# 401202; RRID:AB_2744505 |
| **Biological Samples** |  |  |
| Fresh human palatine tonsil (donor 1) | The University of Chicago Medicine | De-identified donor; IRB# 22-1966 |
| Fresh human palatine tonsil (donor 2) | The University of Chicago Medicine | De-identified donor; IRB# 22-1966 |
| **Chemicals, Peptides, and Recombinant Proteins** |  |  |
| Amicon Ultra-50,000-molecular-weight-cutoff concentrator | EMD Millipore | Cat# UFC5050 |
| Dibenzocyclooctyne-PEG4-N-hydroxysuccinimidyl ester | Sigma-Aldrich | Cat# 764019 |
| Dimethyl sulfoxide | Sigma-Aldrich | Cat# D8418 |
| Blocker BSA | Thermo Scientific | Cat# 37520 |
| Ultrapure salmon sperm DNA solution | Invitrogen | Cat# 15632011 |
| 10 mM ATP | New England Biolabs | Cat# P0756L |
| T4 ligase | New England Biolabs | Cat# M0202L |
| RNase inhibitor, murine | New England Biolabs | Cat# M0314L |
| AMPure XP | Beckman Coulter | Cat# A63881 |
| Buffer EB | Qiagen | Cat# 19086 |
| KAPA HiFi HotStart ReadyMix, 2× | KAPA Biosystems | Cat# KK2601 |
| SPRIselect beads | Beckman Coulter | Cat# B23317 |
| USER Enzyme | New England Biolabs | Cat# M5505L |
| **Critical commercial assays** |  |  |
| Visium Spatial Gene Expression Slide & Reagent Kit, 4 rxns | 10x Genomics | Cat# PN-1000187 |
| Library Construction Kit, 4 rxns | 10x Genomics | Cat# PN-1000196 |
| **Oligonucleotides** |  |  |
| Probes, Sequencing primers, Connector oligomers | IDT | see Table S2 and Table S4 |
| **Deposited data** |  |  |
| Sequencing data | This paper | https://www.ncbi.nlm.nih.gov/geo/query/acc.cgi?acc=GSE304749 |
| **Software and algorithms** |  |  |
| Python code | This paper | https://github.com/jjxia955/Spatial-Prox-seq |

### Experimental Model and Subject Details

#### Human tonsil samples

Fresh human tonsil samples were obtained from The University of Chicago Medicine in accordance with institutional guidelines. This study was approved by The University of Chicago Institutional Review Board (IRB# 22-1966). The samples (de-identified) were first embedded in optimal cutting temperature (OCT) compound. They were then stored in a sealed container at –80°C. Embedded tissue blocks were sectioned into 10-μm slices. They were then mounted on microscope slides (Fisherbrand Superfrost Plus Microscope Slides, 12-550-15) for immunofluorescence staining or on Visium slides (10x Genomics) for Sprox-seq experiments.

### Method details

#### Antibodies used for Sprox-seq

The antibodies and clones were listed below: CD19 (HIB19); CD20 (2H7); CD21 (Bu32); CD35 (9H3); CD23 (EBVCS-5); IgM (MHM-88); IgD (IA6-2); Human IgG (12G8G11); CD32 (FUN-2); CD38 (HB-7); CD24 (ML5); CD3 (HIT3a); CD4 (RPA-T4); CD8 (SK1); CD28 (CD28.2); CD40 (5C3); CD147 (HIM6); CD279 (A17188B); CD27 (O323); CD86 (IT2.2); PDL2 (24F.10C12); CD9 (H19a); CD81 (5A6); ICAM1 (HA58); VCAM1 (P3C4); CD29 (TS2/16); ITGA4 (HP2/1); LFA1 (m24); CD11a (CRIS-3); HLADPRQ (CR3/43); Isotype control mouse IgG1 (mG1-45); mouse IgG2a (mG2a-53); mouse IgG2b (mG2b-57). The detailed information of all these antibodies was shown in Table S1.

#### DNA oligos synthesis

All PLA oligomers were synthesized by Integrated DNA Technologies. The oligomers were purified using high-performance liquid chromatography (HPLC). Oligomers were dissolved in phosphate-buffered saline (PBS) at a concentration of 80 μM and stored at −20°C. The sequences were based on those published previously^53^, with one modification: six UTP nucleotides were added to the 3’ end of PLA oligomer B. This modification enables the USER enzyme (NEB, M5505S) to cleave the ligated products after the ligation step, which improves the capture efficiency. All PLA oligomer sequences are listed in Table S2.

#### Preparation of antibody-oligonucleotide conjugates (AOCs)

AOCs were prepared as described in other published studies^16,53^. Briefly, antibodies were buffer exchanged into PBS using 50K molecular weight cutoff (MWCO) concentrators (EMD Millipore). The antibodies were then diluted to approximately 2 mg/mL. They were incubated on ice for one hour with 2 mM dibenzocyclooctyne-PEG4-N-hydroxysuccinimidyl ester (DBCO; Sigma, 764019) at a 9:1 volume ratio. After incubation, DBCO-conjugated antibodies were purified using 50K MWCO concentrators. The purified antibodies were diluted to approximately 8 μM and mixed with 80 μM azide-functionalized PLA oligomers at a 1:3 volume ratio. This mixture was incubated at 4°C overnight. Unconjugated PLA oligomers were removed by purification using 50K MWCO concentrators. Glycerol was then added to a final concentration of 50% for storage at −20°C.

#### Immunofluorescence staining validation of antibodies and antibody-oligonucleotide conjugates

The specificity of primary antibodies and AOCs was validated using human tonsil tissue sections. Slides containing human tonsil sections were placed on a thermocycler adapter (10x Genomics, 1000194) for 1 minute at 37°C. Slides were then fixed in 1% paraformaldehyde (PFA) for 10 minutes and washed three times with 3× SSC buffer. Slides were then blocked with 1% bovine serum albumin (BSA) in PBS for 10 minutes, and stained with either antibodies or AOCs at 4°C for 1.5 hours. After staining, the slides were washed three times with 1% BSA and incubated with fluorescently labeled secondary antibodies.

#### Sprox-seq on 10x Visium platform

See Supplementary Information for a detailed description of the Sprox-seq workflow on the 10x Visium platform. Briefly, 10-μm-thick tissue sections were mounted on Visium slides. Prior to processing, slides were placed on a 37°C heat block for 1 minute, and fixed in 1% PFA at room temperature for 10 minutes, followed by three washes with 3× SSC buffer. Slides were then placed on a cassette, blocked with blocking buffer at room temperature for 10 minutes, and incubated with 90 µL of Probe Binding Buffer containing AOCs (each at 2.5 nM) at 4°C for 1.5 hours. After incubation, slides were washed four times with Wash Buffer 1 and incubated with 90 µL of Ligation Buffer at 37°C for 30 minutes. Slides were subsequently washed four times with Wash Buffer 1. USER Enzyme Buffer was then added and incubated at 37°C for 30 minutes, followed by additional washes with Wash Buffer 1. After washing, slides were removed from the cassette and dipped into 3× SSC buffer 20 times prior to imaging. Following imaging, slides were remounted, treated with Decrosslinking Buffer (1× SSC) at 70°C for 15 minutes, and equilibrated at room temperature for 10 minutes. After removing 1× SSC, 100 µL of Wash Buffer 2 was added and incubated at room temperature for 1 minute. Wash Buffer 2 was removed, and 70 µL of Permeabilization Buffer was applied dropwise and incubated at 37°C for 25 minutes. Permeabilization Buffer was then removed, and wells were gently washed twice with 100 µL of 0.1× SSC.

Subsequent steps followed the 10x Visium Spatial Gene Expression User Guide, starting from reverse transcription (Step 1.2) through cDNA amplification, with two modifications: during second-strand synthesis and cDNA amplification, U-Fwd primer was added to enhance amplification of PLA products. After cDNA amplification, PLA products and mRNA-derived cDNAs were separated using 0.6× SPRIselect reagent, with the bead fraction containing mRNA-derived cDNAs and the supernatant containing PLA products. For cDNA library preparation, standard Visium instructions were followed. For PLA library preparation, 70 µL of SPRIselect reagent (2.1×) was added to 75 µL of the supernatant to complete PLA product cleanup. Purified PLA products were indexed and amplified using primers (Table S4). Thermal cycling conditions were as follows: 98°C for 2 minutes; 13 cycles of 98°C for 20 seconds, 63°C for 30 seconds, and 72°C for 5 seconds; followed by 72°C for 5 minutes and hold at 4°C. Following amplification, PLA products were purified with 1.2× SPRIselect reagent and eluted in 25.5 µL of Buffer EB. A total of 25 µL of the eluate was transferred to a new tube, and product concentration was measured using TapeStation analysis. Samples were subsequently prepared for sequencing.

#### Fisher’s Exact Test

To filter out proximity noise, a one-sided Fisher’s Exact Test is employed using a 2×2 contingency table that quantifies co-occurrence counts of two probes (Probe A and Probe B) in each spot. Specifically, for a given PLA product O_i,j_ between Probe A = *i* and Probe B = *j*, the table partitions observed counts into four categories: co-binding events between probes *i* and *j* (i.e., O_i,j_), instances where Probe A is *i* but B is not *j* (i.e., Σ_l≠j_O_i,l_), instances where Probe B is *j* but A is not *i* (i.e., Σ_l≠i_O_i,l_), and all remaining combinations (i.e., Σ_k≠i_Σ_l≠j_O_k,l_). The Fisher’s Exact Test then evaluates whether the observed co-binding between *i* and *j* is significantly enriched compared to other combinations, under the null hypothesis of independence. To account for multiple comparisons, Benjamini-Hochberg correction is applied to the p-values obtained for all PLA products within each spot.

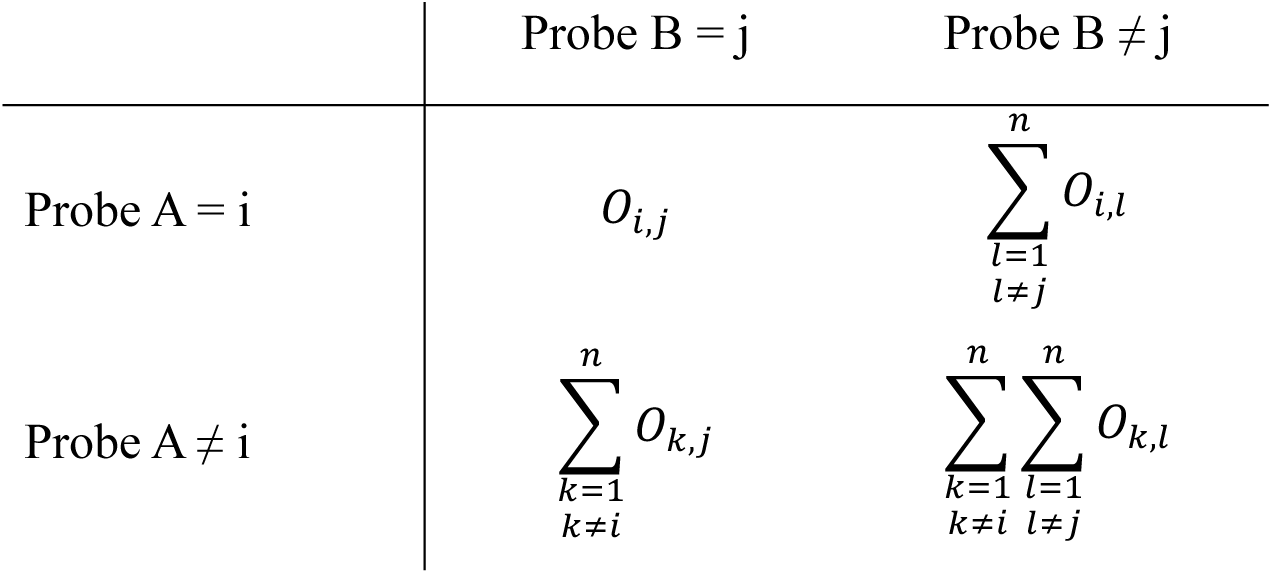

A PLA pair is considered as ‘protein complex’, if the corrected p-value falls below a threshold of 0.05, indicating significant protein interactions beyond random proximity. To quantify the spatial prevalence of the interaction, we define a metric called the **ratio of positive spots**, which reflects the frequency or commonality of the corresponding protein interaction across the entire tissue or within specific regions.

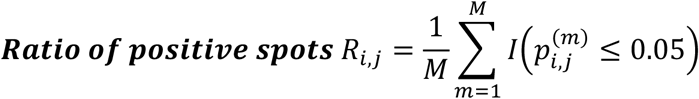

M is the total number of spatial spots evaluated. I is the indicator function, which returns 1 if the interaction is significant in that spot, and 0 otherwise.

#### Protein fractional overlap score

To quantify the relative interaction strength of PLA pairs between two proteins in each spot, denoted as P_i_ and P_j_, we compute two fractional overlap values: F_i_ and F_j_. F_i_ represents the fraction of PLA pair between P_i_ and P_j_ normalized by the total abundance of protein P_i_, while F_j_ does the same relative to protein P_j_. These are defined as:

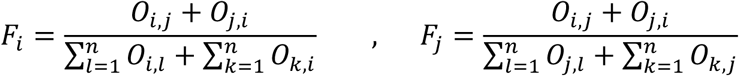

The final protein fractional overlap score between the PLA pair is then computed as:

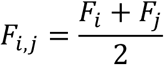

This metric provides an intuitive and normalized measure of the proportion of ligation events shared between two proteins, relative to their total abundance. Protein fractional overlap score is a soft and interpretation-friendly metric that reflects interaction strength in a relative, abundance-aware manner.

#### Next generation sequencing

The cDNA library, with 1% vol/vol PhiX, was sequenced using an Illumina NextSeq 2000 200-cycle P3 kit. The following read distributions were assigned for the cDNA sequencing: read 1, 28 cycles; i7 index, 10 cycles; i5 index, 10 cycles; read 2, 90 cycles. The PLA library, with 40% vol/vol PhiX, was sequenced using an Illumina NextSeq 550 high-output 150-cycle kit.

Custom read 2 primer and custom i7 read primer were prepared and spiked into the sequencing reagent cartridge according to Illumina’s protocol for custom primers (Table S4). The following read distributions were assigned for the PLA sequencing: read 1, 28 cycles; i7 index, 8 cycles; read 2, 75 cycles1.

#### Sequencing alignment

Spatial transcriptomics data generated using the 10x Genomics Visium platform were processed and aligned using Space Ranger 2.0.1. The sequencing data of PLA were aligned with a custom Java program which is deposited at https://github.com/tay-lab/Prox-seq. An alignment guide is also provided for more details. Raw base call (BCL) files from the Illumina sequencer were demultiplexed into FASTQ format using the Illumina BaseSpace platform.

---

**Figure S1.**
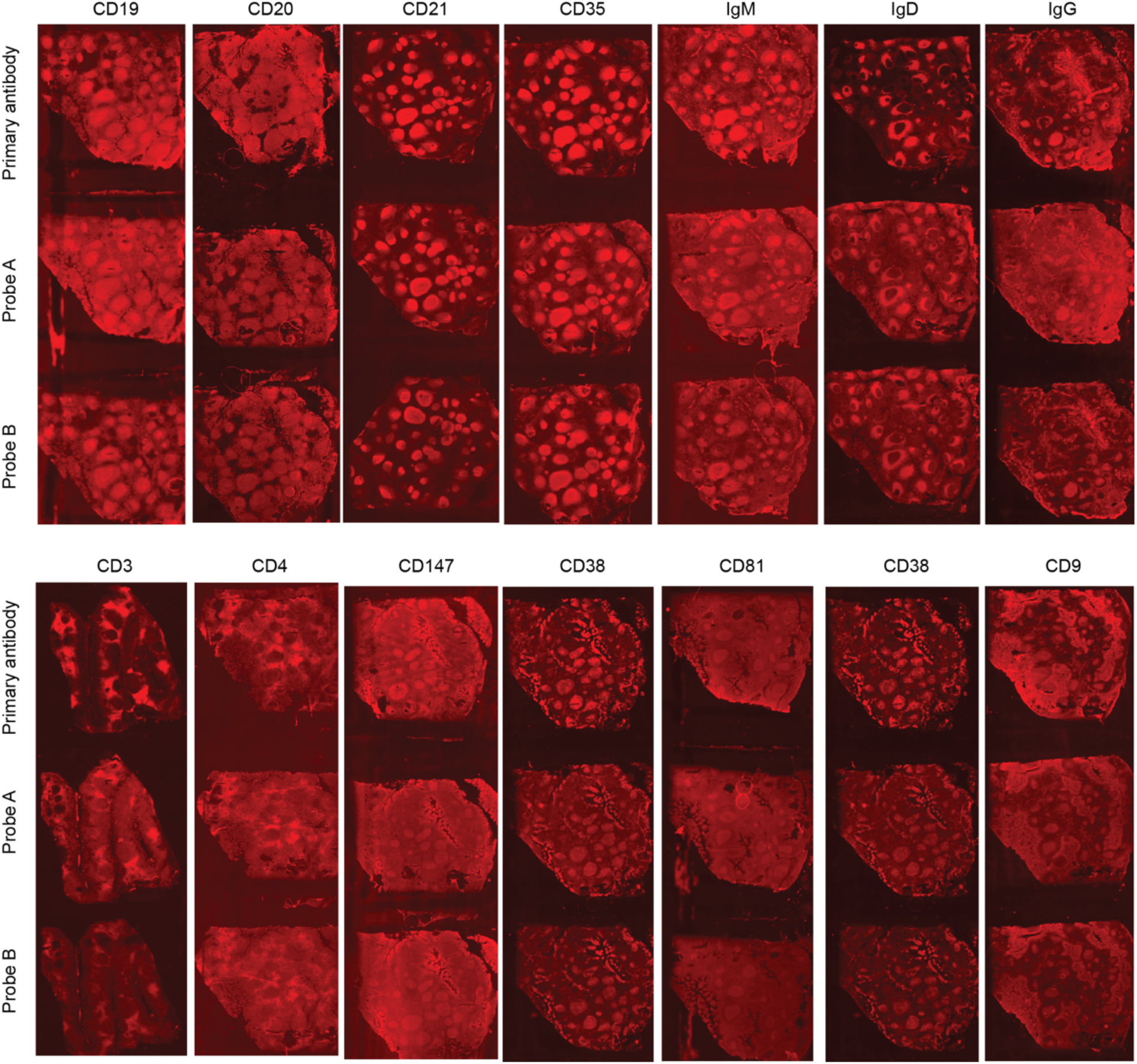
Specificity validation of primary antibodies and PLA probes by immunofluorescence. Related to Figure 1. Specificity of antibodies targeting selected marker proteins was validated by immunostaining for primary antibodies, as well as Probe A and Probe B individually. Representative targets are shown; Validation images of other antibodies are available upon request.

**Figure S2.**
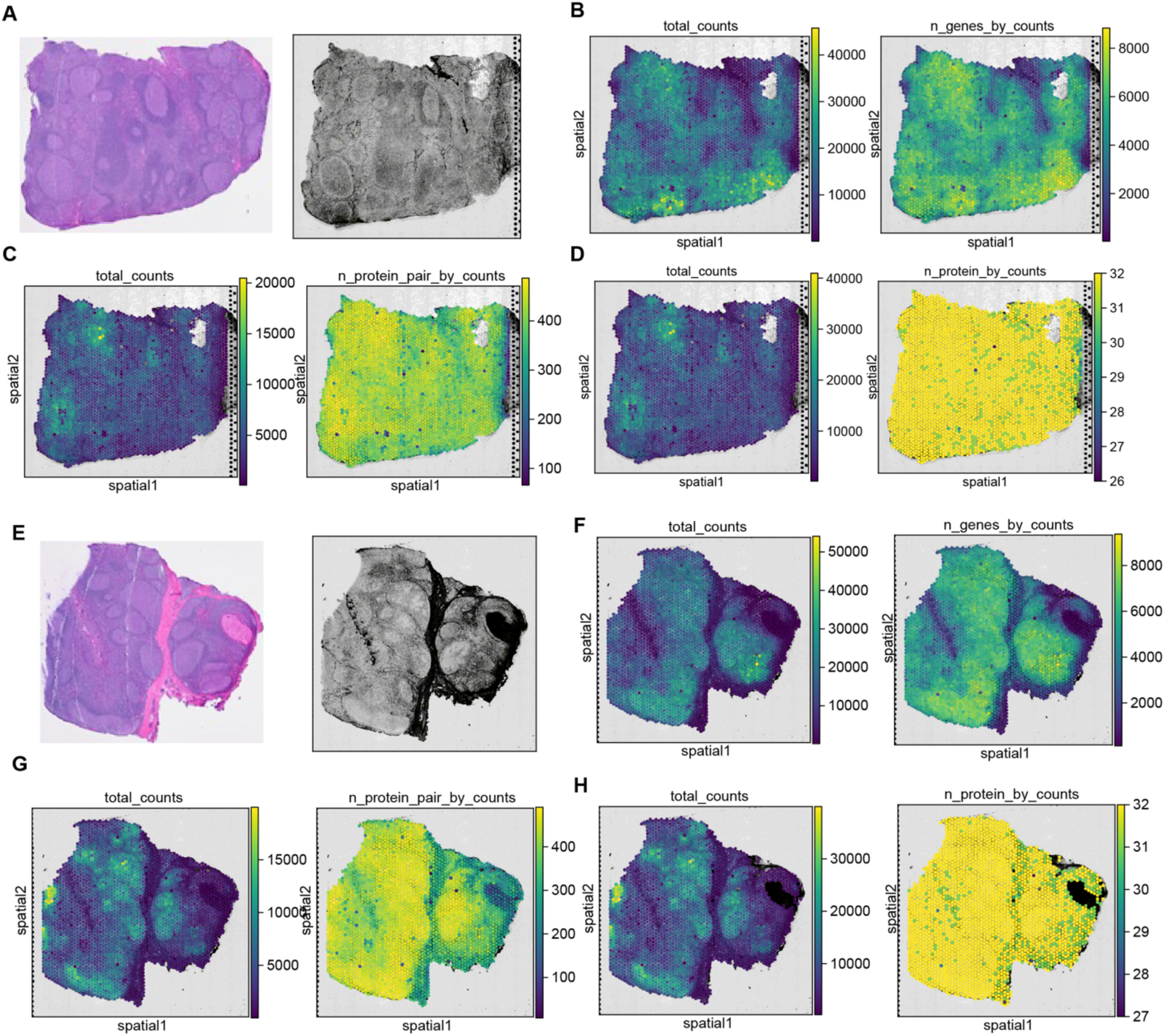
Capture summary of samples B1 and D1. Related to Figure 1. (A) Histological reference and imaging for sample B1. H&E staining of an adjacent tissue section (left) and the brightfield image of the Visium slide used for spatial capture (right). (B– D) Quality measurement data for sample B1. Total mRNA UMI counts and number of detected genes per spot (B). Total PLA UMI counts and number of detected PLA pairs per spot (C). Total protein UMI counts and number of detected proteins per spot (D). € Histological reference and imaging for sample D1. H&E staining of an adjacent tissue section (left) and the brightfield image of the Visium slide used for spatial capture (right). (F– H) Quality measurement data for sample D1. Total mRNA UMI counts and number of detected genes per spot (F). Total PLA UMI counts and number of detected PLA pairs per spot (G). Total protein UMI counts and number of detected proteins (derived from PLA data) per spot (H).

**Figure S3.**
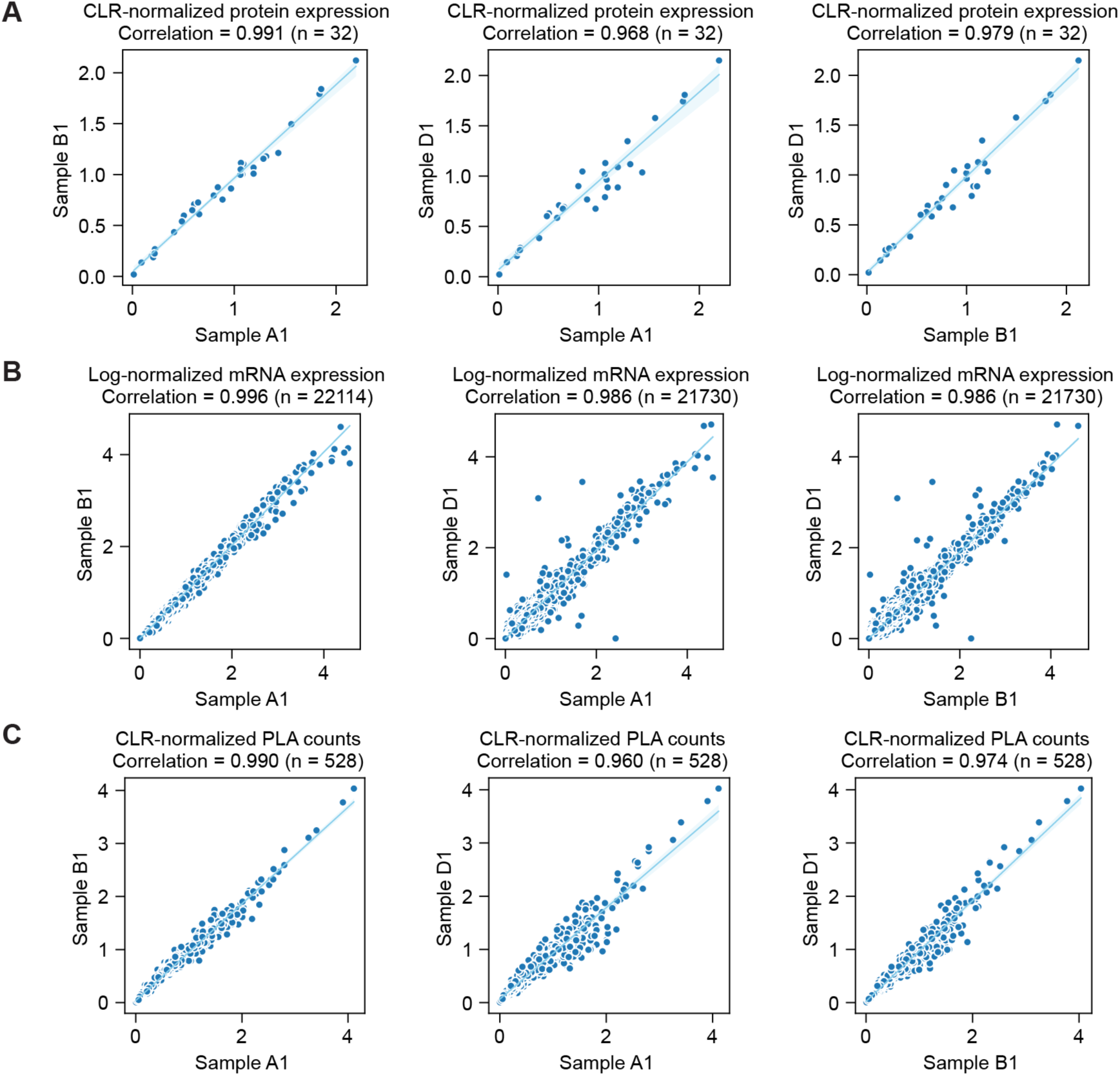
Pearson correlation coefficient analysis across biological replicates. Related to Figure 1. (A–C) Pairwise correlations of normalized mean protein expression (A), mRNA expression (B) and PLA counts (C) across three biological replicates.

**Figure S4.**
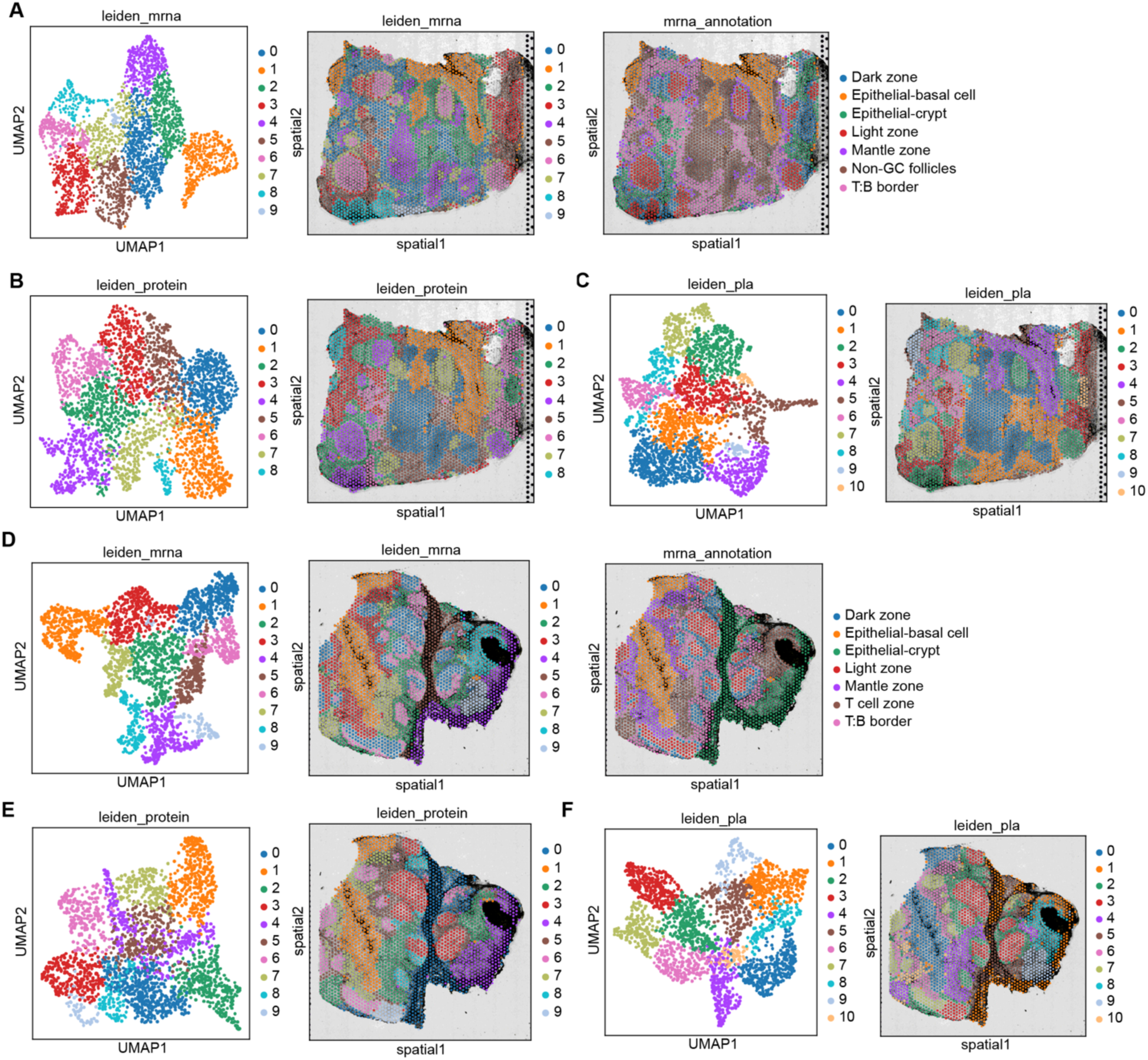
Clustering analysis for samples B1 and D1. Related to Figure 1. (A–C) Clustering analysis for sample B1. (A) Clustering based on RNA data: UMAP plot (left), spatial distribution (middle), and cluster annotation based on H&E morphology and marker gene expression (right). (B) Clustering based on normalized protein data: UMAP plot (left) and spatial distribution (right). (C) Clustering based on normalized PLA data: UMAP plot (left) and spatial distribution (right). (D–F) Clustering analysis for sample D1. (D) Clustering based on RNA data: UMAP plot (left), spatial distribution (middle), and cluster annotation based on H&E morphology and gene expression (right). (E) Clustering based on normalized protein data: UMAP plot (left) and spatial distribution (right). (F) Clustering based on normalized PLA data: UMAP plot (left) and spatial distribution (right).

**Figure S5.**
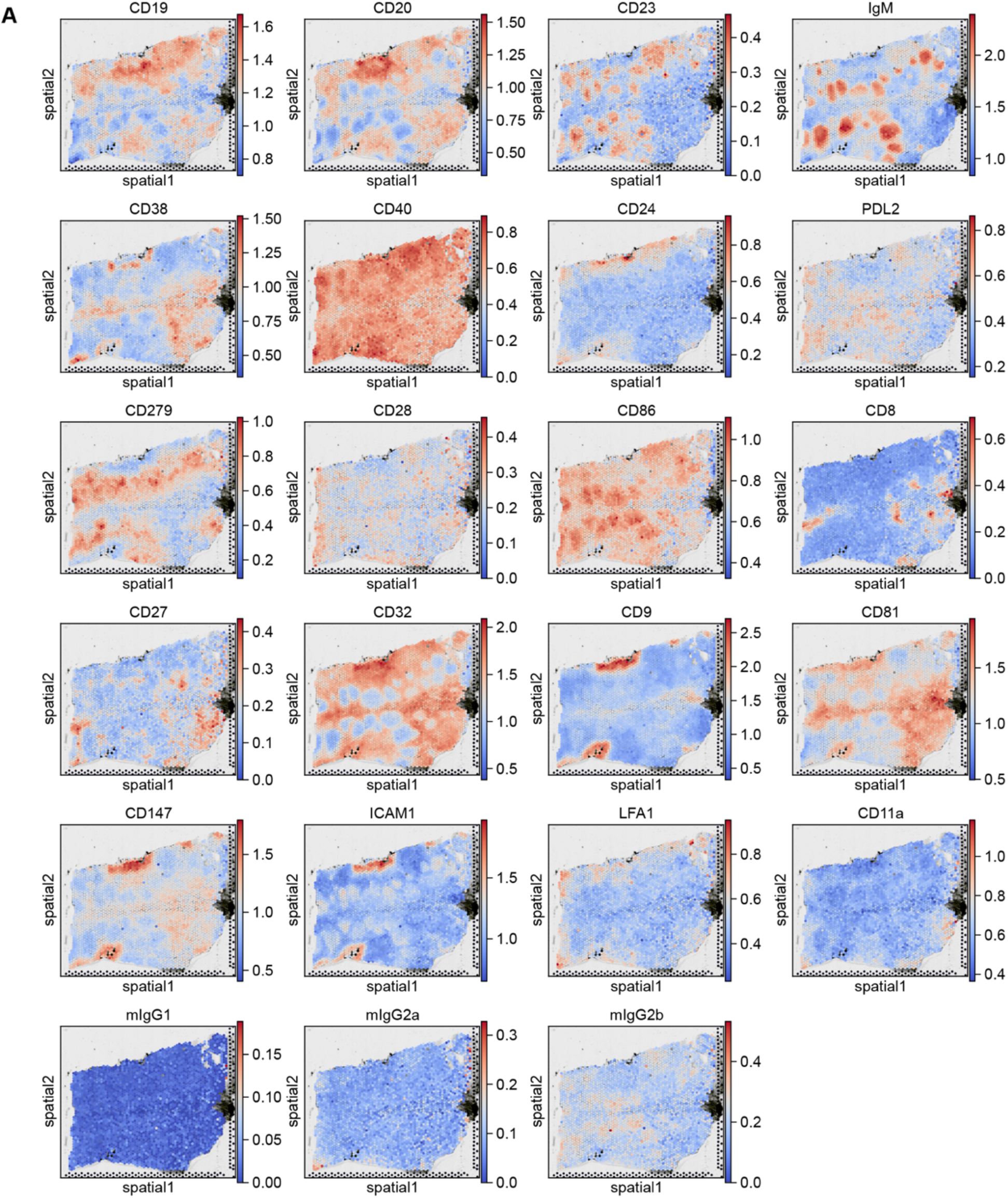
Spatial distribution of additional proteins in the panel for sample A1. Related to Figure 2. (A) Spatial plots showing the expression patterns of the remaining proteins in the panel for sample A1. Normalized protein expression levels are used. ITGA4, CD29 and VCAM1 are shown in Figure 6.

**Figure S6.**
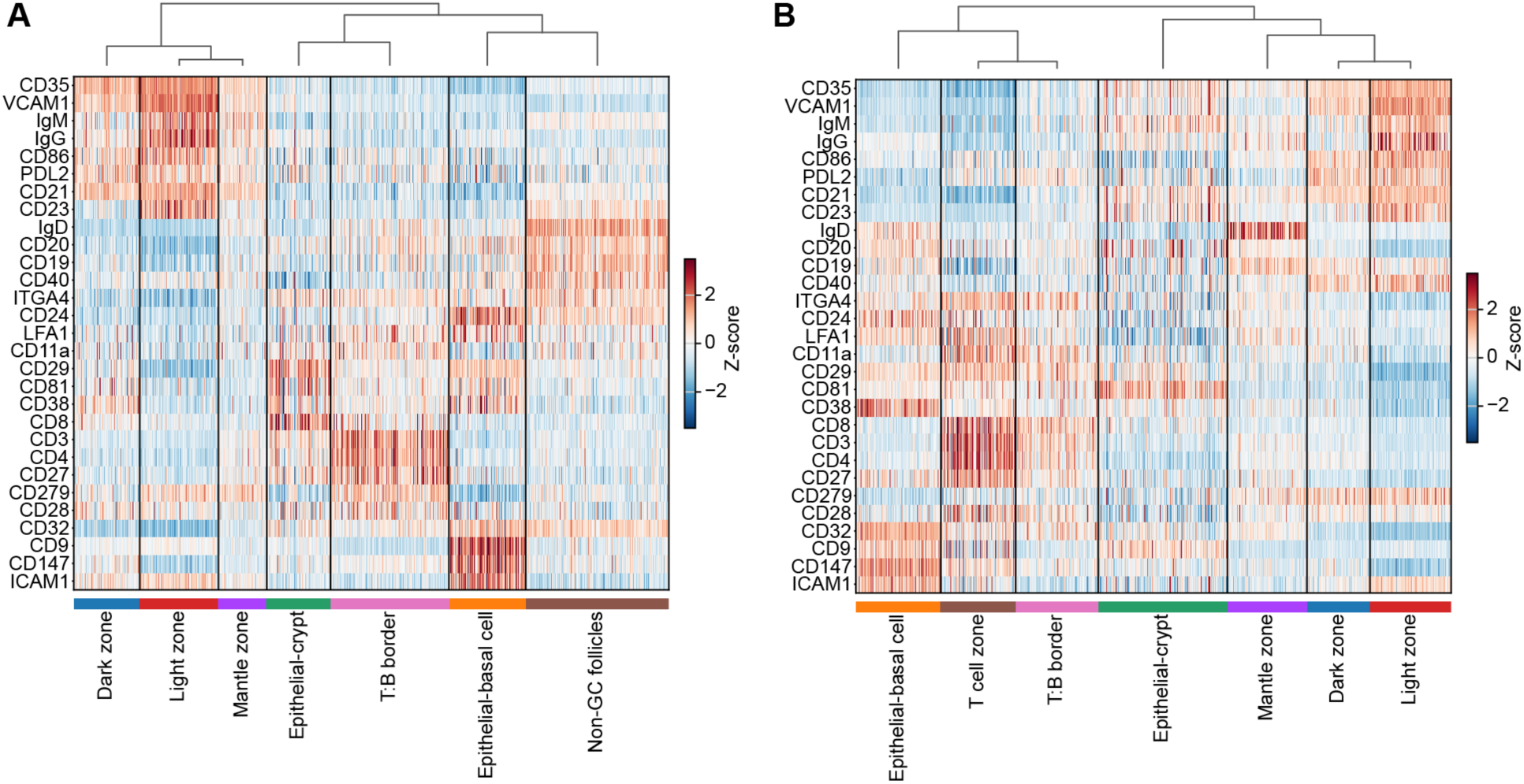
Protein expression patterns across spatial clusters in samples B1 and D1. Related to Figure 2. (A–B) Heatmaps showing protein expression across spatial regions for biological replicates sample B1 (A) and sample D1 (B). Rows represent individual proteins, and columns representing spots are grouped by clusters annotated based on RNA. Normalized expression values were scaled to visualize relative protein abundance across regions.

**Figure S7.**
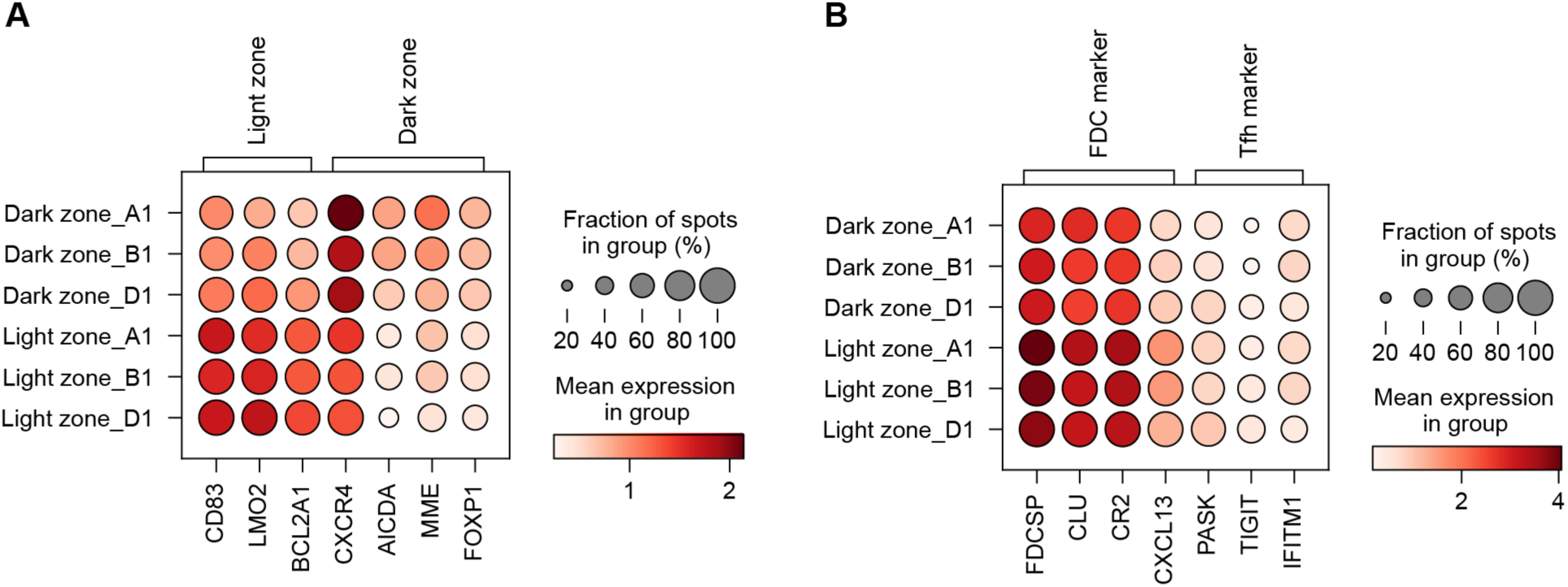
Heatmap of representative markers for Light zone and Dark zone. Related to Figure 2. (A–B) Dot plots showing representative marker expression for Light zone (CD83, LMO2, BCL2A1) and Dark zone (CXCR4, AICDA, MME, FOXP1) (A), as well as follicular dendritic cell (FDC) (FDCSP, CLU, CR2, CXCL13) and T follicular helper (Tfh) cell ( PASK, TIGIT, IFITM1) (B) across three biological replicates. Dot size represents the proportion of spatial spots within each group, and color indicates the mean log-normalized expression level.

**Figure S8.**
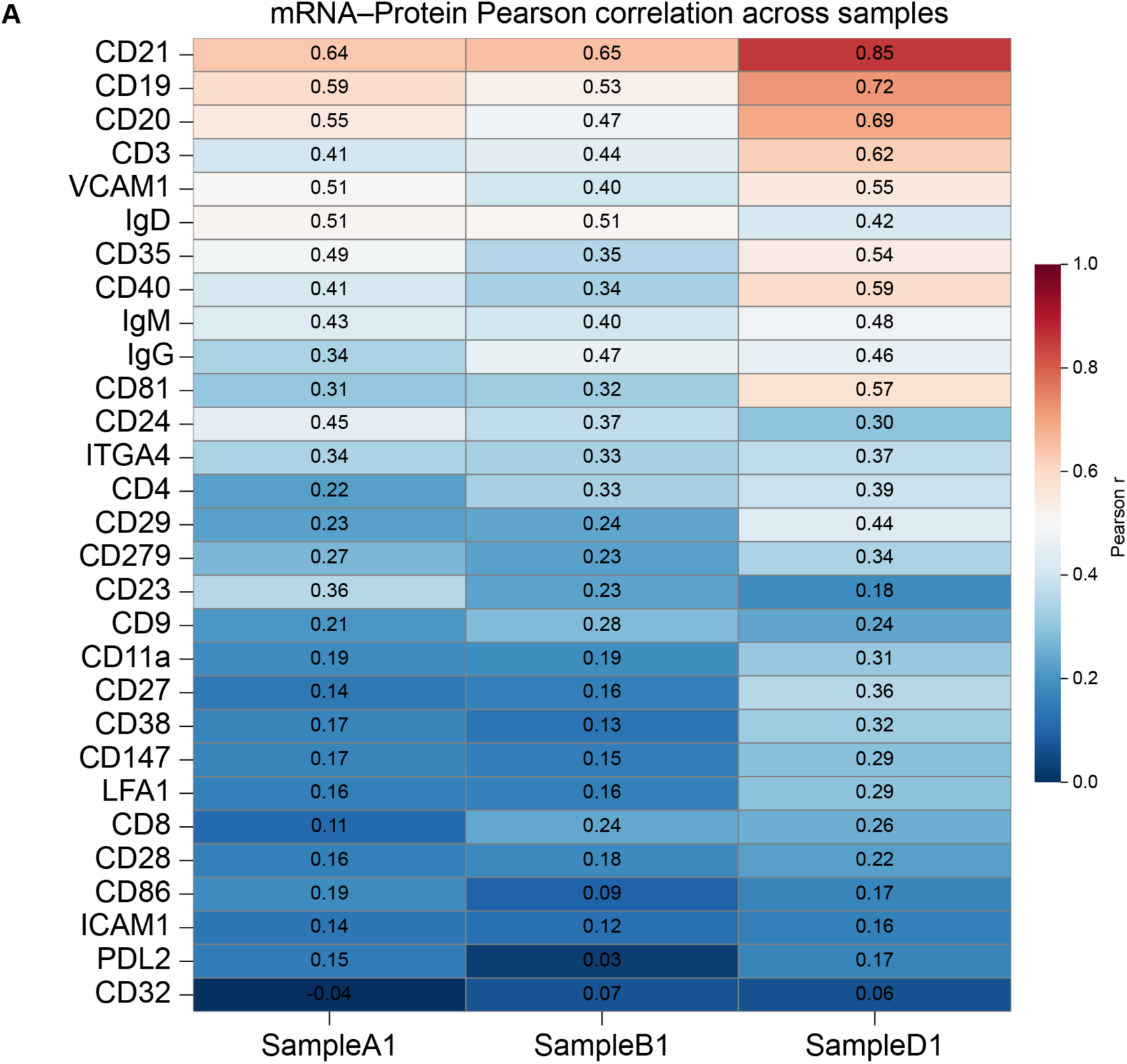
Correlation between mRNA and the protein expression levels. Related to Figure 2. (A) Heatmap showing the Pearson correlation coefficients between mRNA and protein expression levels across three biological replicates. Correlations were calculated using raw counts. The color scale indicates the magnitude of the correlation (r).

**Figure S9.**
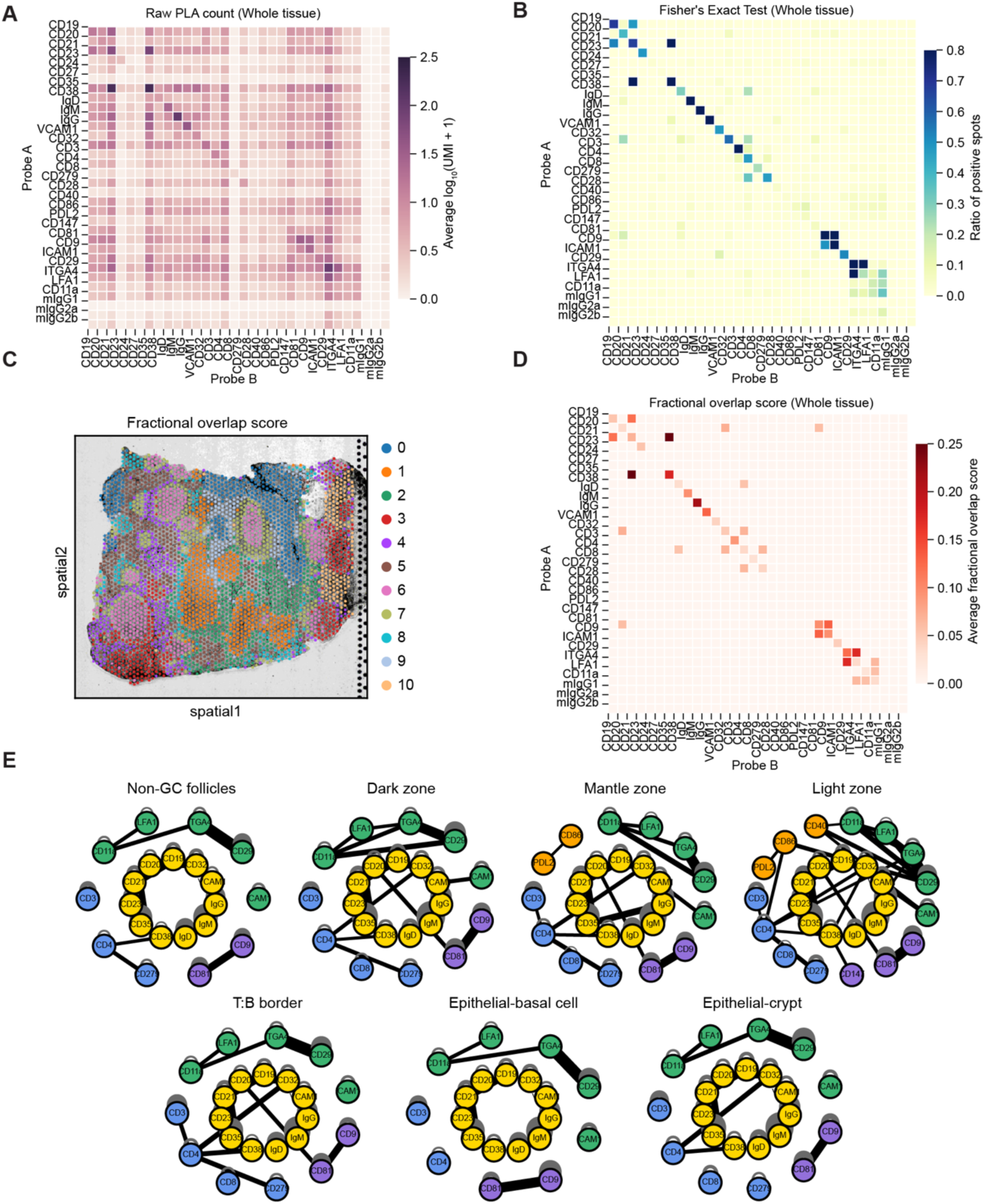
Protein interaction networks in sample B1. Related to Figure 3. (A) Heatmap showing raw PLA counts (log-transformed) for all detected PLA pairs across sample B1. (B) Heatmap of statistically significant interactions identified by Fisher’s Exact Test (adjusted *P* < 0.05) across sample B1. Values represent the ratio of positive spots. (C) Spatial distributions of clustering analysis based on protein fraction overlap scores of protein pairs. (D) Heatmap of protein fractional overlap scores for protein pairs across sample B1. Only protein pairs with a ratio of positive spots greater than 0.05 are shown. (E) Protein interaction networks constructed for each spatial cluster of sample B1.

**Figure S10.**
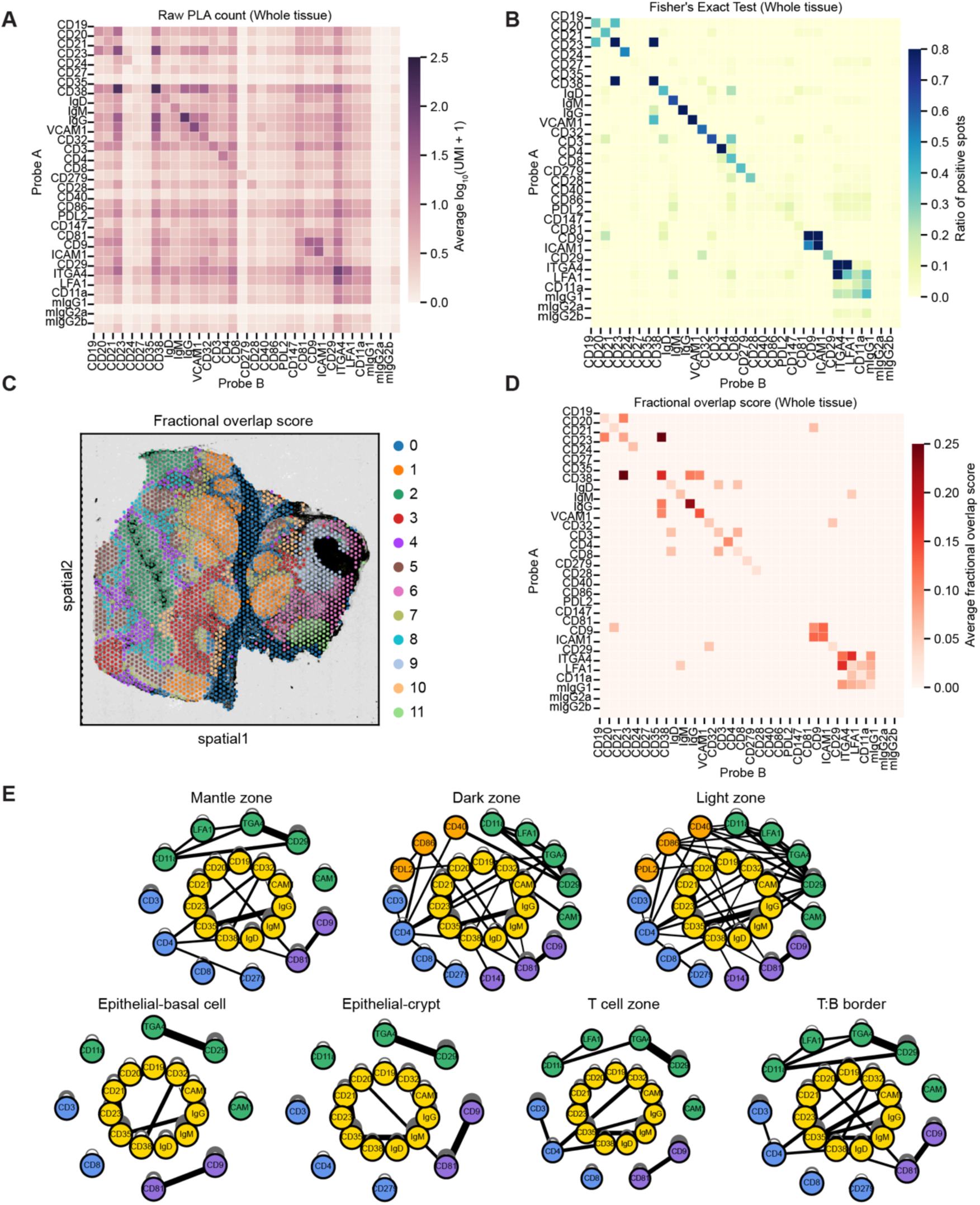
Protein interaction networks in sample D1. Related to Figure 3. (A) Heatmap showing raw PLA counts (log-transformed) for all detected PLA pairs across sample D1. (B) Heatmap of statistically significant interactions identified by Fisher’s Exact Test (adjusted *P* < 0.05) across sample D1. Values represent the ratio of positive spots. (C) Spatial distributions of clustering analysis based on protein fraction overlap scores of protein pairs. (D) Heatmap of protein fractional overlap scores for protein pairs across sample D1. Only protein pairs with a ratio of positive spots greater than 0.05 are shown. (E) Protein interaction networks constructed for each spatial cluster of sample D1.

**Figure S11.**
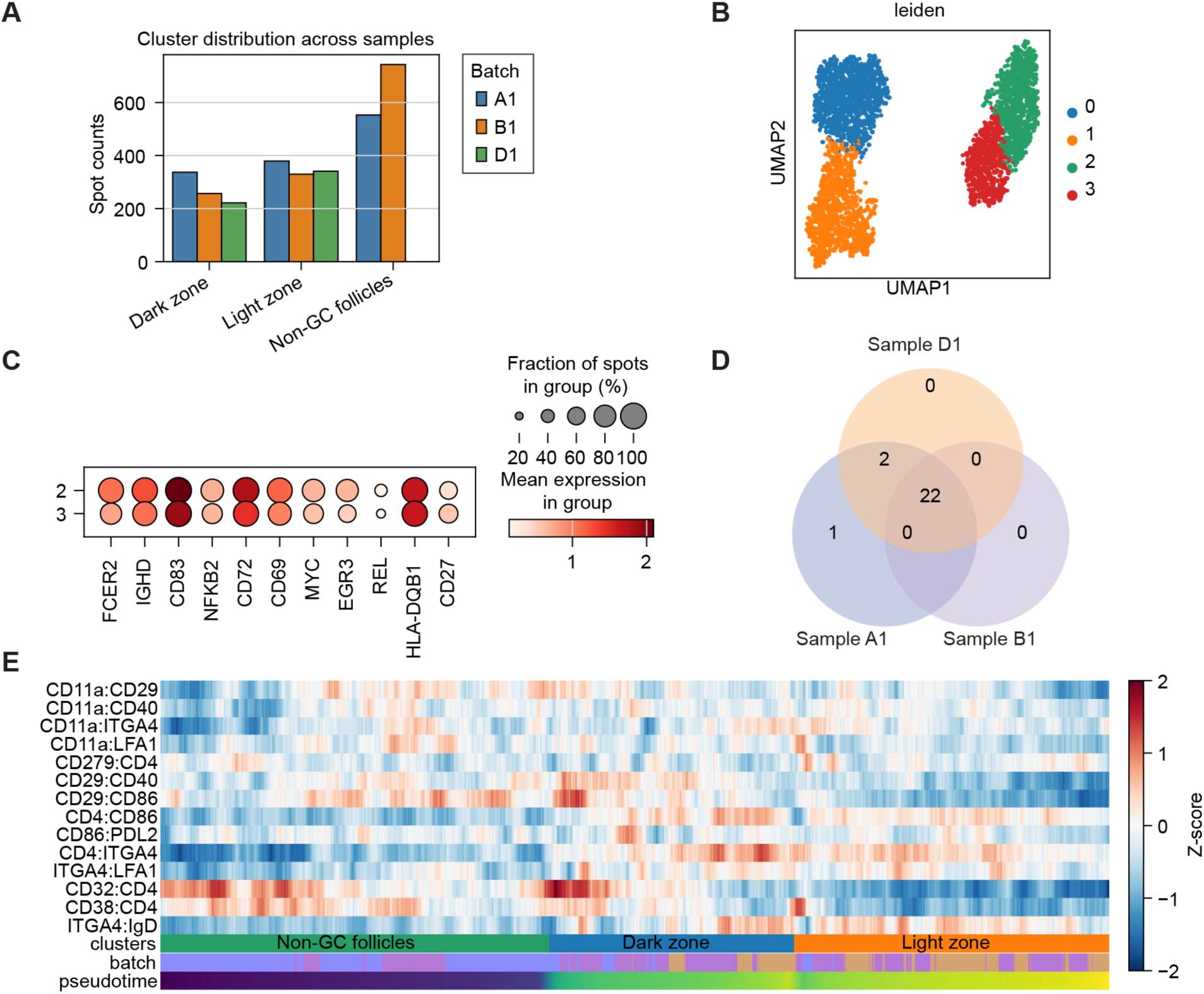
Pseudotime analysis. Related to Figure 4. (A) Bar plot showing sample distribution of selected spots used for pseudotime trajectory inference. (B) UMAP plot showing Leiden clustering of spots based on RNA expression data from three biological replicates, Non-GC follicles are separated into two clusters: Cluster 2 and Cluster 3. (C) Dot plot comparing the expression level of B cell active marker genes between Cluster 2 and 3, indicating a higher activation state in Cluster 2. (D) Venn diagrams showing the number of consistently detected homodimers across three biological replicates. Protein pairs were defined as consistent if they passed Fisher’s Exact Test in all three replicates and exhibited a positive spot ratio greater than 0.15 in at least one of the selected clusters within each sample. (E) Heatmap showing scaled protein fractional overlap scores for integrin molecule-related protein pairs along the pseudotime trajectory. Spots are ordered based on pseudotime values and grouped by clusters. Only consistent protein pairs are included. Z-score normalization was applied to each protein pair across selected spots.

**Figure S12.**
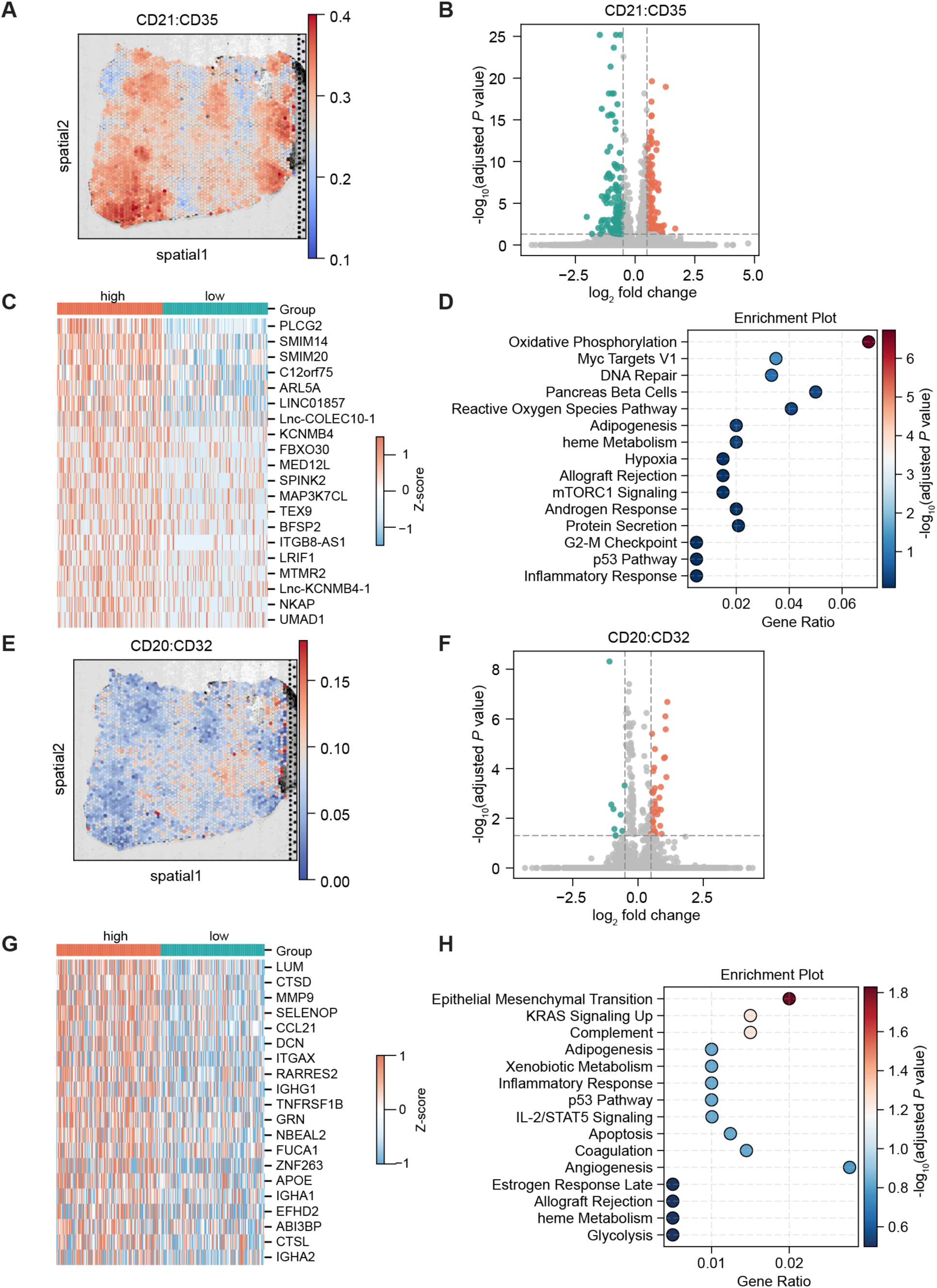
Sprox-seq enables the functional annotation of spatially enriched protein complexes. Related to Figure 5. (A) Spatial plot of CD21 – CD35 raw protein fractional overlap scores in sample B1. (B) Volcano plot of differentially expressed genes between high and low CD21 – CD35 interaction spots within Germinal Centers (Light zone and Dark zone) in sample B1. High and low groups were defined as the top 20% and bottom 20% of protein fractional overlap scores. (C) Heatmap showing the top 20 upregulated genes. (D) Hallmark pathway enrichment analysis (MSigDB Hallmark 2020) of 187 upregulated genes in high CD21–CD35 interaction group. (E) Spatial plot of CD20–CD32 raw protein fractional overlap scores in sample A1. (F) Volcano plot of differentially expressed genes between high and low CD20–CD32 spots within Non-GC follicles from samples A1 and B1. High and low groups were defined as the top 20% and bottom 20% of protein fractional overlap scores. (G) Heatmap showing the top 20 upregulated genes. (H) Hallmark pathway enrichment analysis (MSigDB Hallmark 2020) of 38 upregulated genes in high CD20–CD32 interaction group. Gene differential expression analsyis for panels B and F was analyzed using the Wilcoxon rank-sum test and adjusted *P* values were calculated by the Benjamini–Hochberg method. Upregulated genes (log_2_(fold change) > 0.5 and adjusted *P* < 0.05) are labeled as orange; Downregulated genes (log_2_(fold change) < -0.5 and adjusted *P* < 0.05) are labeled as teal.

**Figure S13.**
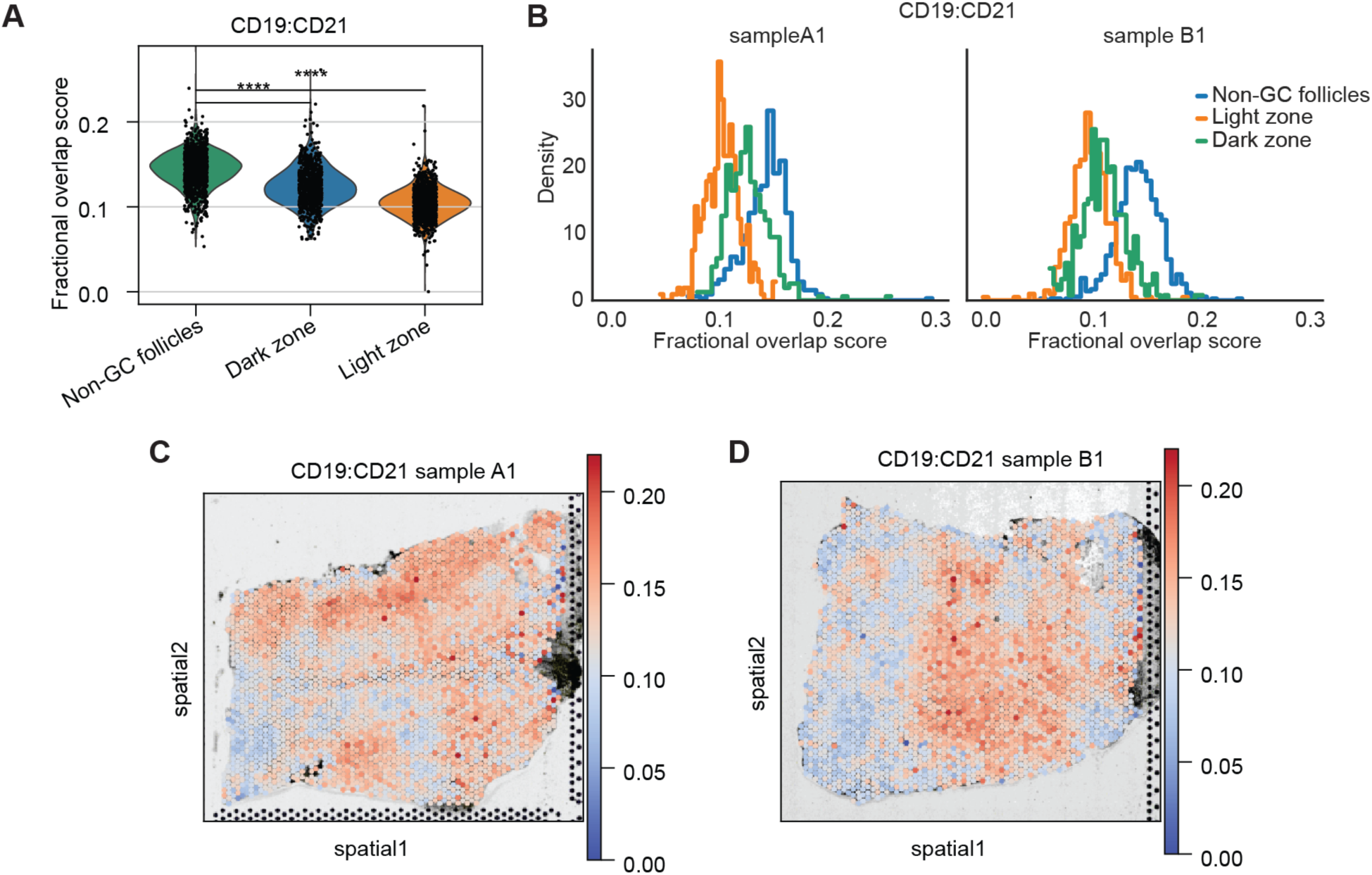
CD19–CD21 interaction in Non-GC follicles. Related to Figure 5. (A) Violin plot of raw protein fractional overlap scores for CD19– CD21 across selected clusters of three samples. Each dot represents a single spatial spot. (B) Histograms of raw protein fractional overlap scores of CD19–CD21 across tissue spots from samples A1 and B1, grouped by clusters. (C–D) Spatial plot of raw protein fractional overlap scores for CD19–CD21 in samples A1 (C) and B1 (D). Statistical significance of panel A was analyzed using the two-sided Mann–Whitney U test; asterisks indicate significance levels: *P* < 0.0001 (****).

**Figure S14.**
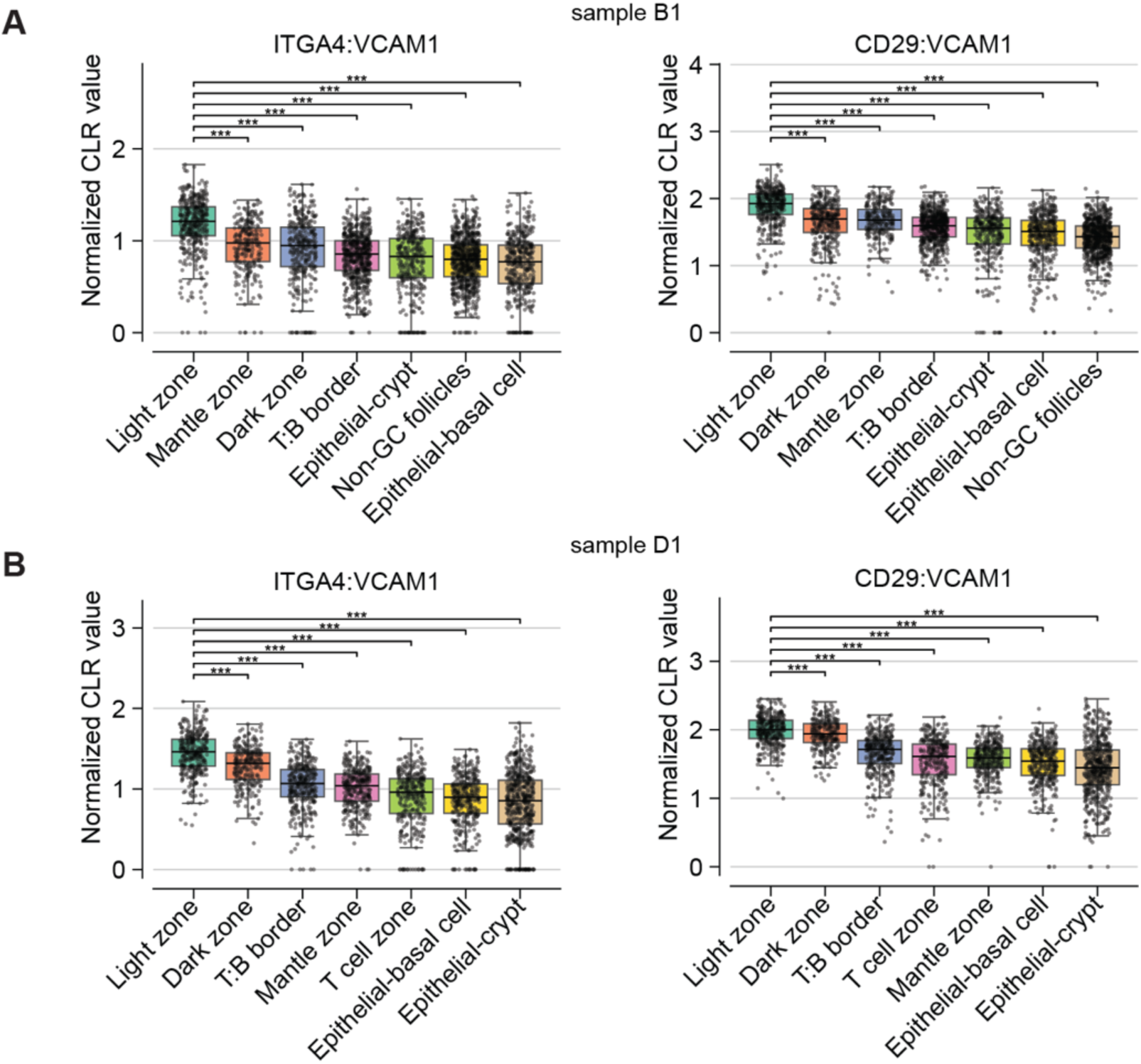
Comparison of ITGA4–VCAM1 and CD29–VCAM1 interaction in clusters across samples B1 and D1. Related to Figure 6. (A–B) Comparison of normalized value of ITGA4–VCAM1 (left) and CD29–VCAM1 (right) across all clusters in samples B1 (A) and D1 (B). Statistical significance for panels A and B was assessed using the two-sided Mann–Whitney U test and *P* values were adjusted by the Benjamini–Hochberg method; asterisks indicate significance levels: *P* < 0.001 (***).

## Notes

https://www.ncbi.nlm.nih.gov/geo/query/acc.cgi?acc=GSE304749

https://github.com/jjxia955/Spatial-Prox-seq

### Competing Interest Statement

The authors have declared no competing interest.

