## supplementary information for "Spatial Proximity Sequencing Maps Developmental Dynamics in the Germinal Center"

### Spatial prox-seq on 10x Visium platform

**This protocol is used with the Visium Spatial Gene Expression Reagent Kits from10x Genomics.**

**Sectioning**

Tissue sectioning was performed following the guidance of Methanol Fixation, H&E Staining & Imaging for Visium Spatial Protocols (10x Genomics). Briefly, OCT-embedded tissue blocks were sectioned into 10-μm slices and mounted on Visium Slides and stored at -80˚C before use.

**Fixation**

(1) Put the slide on a heat block (preheated at 37˚C) for 1 minute.

(2) Transfer the slide into a 50 mL Falcon tube containing 1% PFA (freshly prepared with PBS) and fix for 10 minutes.

(3) Use the Kimwipe to gently remove excess 1% PFA and dip the slide into 3x SSC for four times.

**Staining with antibody-oligonucleotide conjugates and performing proximity ligation**

(1) Gently mount the slide onto the Visium Slide Cassette.

(2) Add 100 μL of Blocking Buffer (1% BSA in PBS, 1U/uL RNase Inhibitor (NEB, M0314L)) to each well. Incubate at room temperature for 10 minutes.

(3) Carefully remove Blocking Buffer from wells by slow pipetting.

(4) Add 90 µL Probe Binding Buffer (0.2% BSA, 0.2 mg/mL sonicated salmon sperm DNA, 100 µg/mL mouse isotype antibodies, 1U/uL RNase Inhibitor) containing antibody-oligonucleotide conjugates (2.5 nM each) to each well. Incubate at 4˚C for 90 minutes.

(5) Wash the wells four times with 100 µL of Wash Buffer-1 (1% BSA in PBS, 1U/uL RNase Inhibitor) at room temperature.

(6) Add 90 µL Ligation Buffer (50 mM HEPES, pH 7.5, 10 mM MgCl2, 1 mM rATP (NEB, P0756S), 9.5 nM connector oligomer (see Table S4), 130 U/mL T4 ligase (NEB), 1U/uL RNase Inhibitor) to each well. Incubate at 37˚C for 30 minutes.

(7) Wash the wells four times with 100 µL of Wash Buffer-1 at room temperature.

(8) Add 50 µL USER Enzyme Buffer (0.1 U/ uL USER, 1x rCutSmart™ Buffer, 1U/uL RNase Inhibitor) to each well. Incubate at 37˚C for 30 minutes.

(9) Wash the wells four times with 100 µL of Wash Buffer-1 at room temperature.

(10) Unmount the slide from the slide cassette and dip it 20 times in a 50 mL Falcon tube filled with 3x SSC buffer. Remove excess liquid from the back of the slide.

(11) Proceed to imaging.

**De-crosslinking PFA-fixed RNA**

(1) After imaging, place the slide back to a slide cassette. Add 100 µL of Decrosslinking Buffer (1x SSC) to each well. Seal the cassette with sealing tape, close the heated lid, and incubate at 70°C for 15 minutes.

(2) Transfer the slide cassette to the bench, gently remove the sealing tape, and equilibrate at room temperature for 10 minutes.

(3) Remove 1x SSC, then add 100 μL of Wash Buffer-2 (0.1% BSA, 0.1% Triton X-100, 3x SSC, 1U/uL RNase Inhibitor) to each well. Incubate at room temperature for 1 minute.

**Permeabilization and reverse transcription**

(1) Preheat the Thermocycler adapter on Thermomixer to 37 ° C.

(2) Remove Wash Buffer-2 from each well.

(3) Add 70 μL of Permeabilization Buffer (3.5 μL Tissue Removal Enzyme (10x Genomics, 3000387), 59.5 μL 1x SSC, 7.0 μL 10% SDS) drop by drop onto the tissue section. Apply sealing tap to the slide cassette and place the cassette on the Thermocycler Adaptor. Close the lid and incubate for 25 minutes.

(3) After incubation, carefully remove the Permeabilization Buffer by slowly pipetting from the corner of the well.

(4) Wash each well twice with 100 μL of 0.1x SSC, pipetting slowly to avoid damaging the tissue.

(4) Proceed with the Visium Spatial Gene Expression User Guide starting from Step 1.2 (Reverse Transcription) to Second Strand Synthesis.

**Second strand synthesis**

(1) For second strand synthesis, add 2 μL of U-fwd (100 uM) primer (see Table S4) to the reaction to amplify the PLA products.

(2) Continue following the User Guide instructions until Step 3.0 (cDNA amplification).

**cDNA amplification and cleanup**

(1) Determine the cDNA PCR cycle number using qPCR, following the User Guide.

(2) After qPCR, prepare the cDNA amplification mix on ice as detailed below, following Step 3.2 of the Visium Spatial Gene Expression User Guide. Include the U-fwd primer to enhance the yield of PLA products.

| cDNA amplification reaction mix | PN | 1x (µL) | 4x+10% (µL) |
|---|---|---|---|
| Amp Mix | 2000047/2000103 | 50 | 220 |
| cDNA primers | 2000089 | 15 | 66 |
| U-Fwd primer (2 µM) | | 2 | 8.8 |

(3) Add 67 µL of the cDNA Amplification Reaction Mix to 35 µL of the sample obtained from second strand synthesis.

(4) Mix thoroughly by pipetting. Briefly centrifuge the mixture.

(5) Continue with cDNA amplification as outlined in Step 3.2 of the Visium Spatial Gene Expression User Guide (step 3.2).

**cDNA and PLA-product size selection**

(1) After cDNA amplification, separate PLA products and mRNA-derived cDNAs using 0.6x SPRIselect reagent. The bead fraction contains mRNA-derived cDNAs and the supernatant contains PLA products. Add 60 µL SPRIselect reagent (0.6x) to cDNA reaction.

(2) Incubate at room temperature for 5 minutes.

(3) Place the tube on the magnet and wait for ~1 minute until the solution is clear.

(4) Carefully transfer and save 75 μL of the supernatant into a new tube strip.

(5) Proceed with cDNA cleanup and library preparation as outlined in Step 3.3 of the Visium Spatial Gene Expression User Guide, continuing through to Step 4.6 to complete cDNA library construction.

**PLA products cleanup**

(1) Vortex to resuspend the SPRIselect reagent. Add 70 μL of SPRIselect reagent (2.1x) to 75 μL of the transferred supernatant and mix by pipetting 15 times.

(2) Incubate the mixture at room temperature for 5 minutes.

(3) Place the tube on the magnet and wait until the solution is clear.

(4) Carefully remove the supernatant.

(5) Add 200 μL of 80% ethanol to the pellet. Wait for 30 sec.

(6) Remove the ethanol.

(7) Repeat steps (5) and (6) for a total of two washes.

(8) Briefly centrifuge and place the tube back on the magnet.

(9) Remove any remaining ethanol and air dry the pellet for 2 minutes.

(10) Remove the tube from the magnet and add 25.5 μL of Buffer EB (Qiagen, 19086). Mix thoroughly by pipetting 15 times.

(11) Incubate at room temperature for 2 minutes.

(12) Place the tube strip back on the magnet until the solution clears.

(13) Transfer 25 μL of the elution to a new tube strip.

(14) Store the elution at 4°C for up to 72 hours, or at −20°C for up to 4 weeks.

**PLA products library amplification**

(1) Use 5 μL of eluted PLA products to set up a 50 μL PCR reaction (1x KAPA HiFi HotStart Readymix, 0.2 μM 10X_SI_PCR primer and 0.2 μM P7_N70X_Custom2 primer (See Table S4)).

(2) Perform thermal cycling with the program: 98°C 2 minutes; 13 cycles of 98°C 20 seconds, 63°C 30 seconds, 72°C 5 seconds; 72°C 5 minutes; 4°C hold.

**PLA products purification**

(1) Vortex the SPRIselect reagent to resuspend. Add 60 μL of SPRIselect Reagent (1.2x) to 50uL of the PCR amplification product. Mix thoroughly by pipetting 15 times.

(2) Incubate at room temperature for 5 minutes.

(3) Place the tube on the magnet and wait until the solution clears.

(4) Carefully remove the supernatant.

(5) Add 200 μL of 80% ethanol to the pellet and wait for 30 seconds.

(6) Remove the ethanol.

(7) Add 200 μL of 80% ethanol to the pellet. Wait for 30 sec.

(8) Remove the ethanol.

(9) Briefly centrifuge the tube and place it back on the magnet. Remove any remaining ethanol.

(10) Take the tube off the magnet and add 25.5 μL Buffer EB. Mix thoroughly by pipetting 15 times.

(11) Incubate at room temperature for 2 minutes.

(12) Place the tube on the magnet until the solution clears.

(13) Transfer 25 μL of the supernatant to a new tube strip.

(14) Perform TapeStation analysis to measure the concentration.

(15) Proceed to sequencing.
